# Secondary structure distances reveal a new dimension of protein evolution

**DOI:** 10.64898/2026.04.29.721599

**Authors:** Adolfo Bastida, Ana María Muñoz, Marcos Egea-Cortines

## Abstract

Molecular phylogenetics based on primary sequence comparisons has been central to reconstructing protein evolution. However, structural evolution does not necessarily parallel sequence divergence, particularly in proteins combining ordered domains with intrinsically disordered regions (IDRs). Here, we introduce a quantitative secondary structure distance (S2D) metric that enables systematic comparison of protein secondary structure, including both ordered elements and IDRs. Using the MADS-box transcription factor family as a model, we show that structural divergence is domain-specific and only partially coupled to sequence-based phylogeny. Domain-resolved analyses reveal that the DNA-binding M domain remains structurally constrained, whereas the I and C domains exhibit extensive sequence divergence while retaining conserved intrinsic disorder. In contrast, the K domain contributes disproportionately to global structural variability. Integrating S2D with phylogenetic distance uncovers both convergent structural architectures among distantly related proteins and pronounced structural remodelling within closely related paralogs—patterns not evident from primary sequence comparisons alone. Residue-level analyses further demonstrate that the structural impact of mutation depends strongly on amino acid identity and does not scale directly with substitution frequency or conservation metrics. Together, these findings indicate that secondary structural evolution provides an additional dimension of protein diversification beyond sequence divergence. By integrating phylogenetic and structural distances, this framework offers a complementary approach to interpreting protein evolution, particularly in families containing mixtures of ordered domains and intrinsically disordered regions.

**Significance Statement:** Evolutionary relationships are typically inferred from primary sequence comparisons, yet structural evolution may follow different trajectories. By developing a quantitative measure of secondary structural divergence, we show that structural change within the MADS-box transcription factor family can both converge and diverge independently of sequence-based phylogeny. Intrinsically disordered regions exhibit extensive sequence divergence while retaining conserved disorder, whereas specific amino acid substitutions disproportionately reshape secondary structure. These findings demonstrate that evolutionary diversification operates through domain-specific structural modulation rather than uniform sequence divergence. Integrating structural and phylogenetic distances provides a complementary framework for interpreting protein evolution and reveals evolutionary patterns that remain hidden when relying on sequence comparisons alone.

## Introduction

The primary structure of proteins constitutes a central source of biological information, enabling the inference of evolutionary relationships, conserved domains, and functional homology. Molecular phylogeny relies on quantifying distances between protein sequences (1). Primary structures are compared using amino acid substitution matrices such as BLOSUM (2). Evolutionary relationships are reconstructed using algorithms including Neighbor Joining, Maximum Likelihood, Parsimony, and Bayesian inference (3–6). A core principle of molecular phylogeny is that mutations accumulate over time. However, at large evolutionary distances, successive substitutions may erode the phylogenetic signal, causing sequences to appear as random amino acid combinations. Loss of phylogenetic signature is often interpreted as reflecting both increased evolutionary distance and functional divergence.

Protein secondary structure (S2) emerges from the primary amino acid sequence. Canonical S2 elements include α-helices, β-sheets, and turns, while intrinsically disordered regions (IDRs), also referred as coils(7), lack stable conformations. Structural classification has expanded to include less frequent motifs such as 3_10_-helices, π-helices and bridges (8). The recognition of IDRs as encoded, functional, and dynamically regulated elements has fundamentally reshaped our understanding of protein structure–function relationships (9). Because IDRs sample conformational ensembles rather than single stable states, their structural characterization requires ensemble-based modeling(10).

The advent of AlphaFold (11) has transformed structural biology by enabling large-scale prediction of protein tertiary structures. This development makes it possible to compare proteins at structural levels beyond primary sequence. Structural alignment tools such as Foldseek(12) and DALI(13) allow 3D comparisons based on structural alphabets or Cα–Cα distance matrices. However, these approaches are optimized for conserved folded cores and are inherently limited in their capacity to characterize IDRs, whose conformations are not fixed but continuously explore conformational space. Consequently, any single set of Cα coordinates represents only one state among many possible configurations. Yet IDRs are central to key biological processes, including phase separation, DNA–protein interactions, and thermosensing (14–16). Current 3D alignment methods therefore overlook or incompletely represent a substantial fraction of functional protein space.

Secondary structure offers an intermediate and potentially informative layer between primary sequence and tertiary folding. Unlike tertiary structures, primary sequence and S2 annotations can be directly aligned using conventional sequence alignment frameworks. This provides an opportunity to incorporate both ordered and disordered regions into structural comparisons.

Here, we introduce a distance metric based on protein secondary structure (S2 distance) designed to quantify similarity while explicitly incorporating IDRs. As a proof of concept, we apply this approach to the MADS-box transcription factor family, which has been extensively characterized phylogenetically and structurally (17, 18). MADS-box proteins are composed of a structured MADS domain, followed by a flexible I domain, a keratin-like K domain, and a largely disordered C-terminal domain. The coexistence of ordered and disordered modules makes this family particularly suitable for testing S2-based comparisons.

Using the S2 distance metric, we identify discrepancies between primary sequence–based phylogenies and S2-derived relationships. Integration of phylogenetic and S2 distances reveals patterns consistent with convergent and divergent structural evolution. Residue-level comparisons further allow prediction of substitutions with strong impact on S2 context. Together, our framework extends evolutionary analysis to the secondary structural level and provides a complementary dimension for detecting structural conservation and divergence across protein families.

## Results

### The MADS-box proteins show staggered ordered-disordered domains

We used a total of 96 MADS-box proteins from Arabidopsis, *Vitis vinifera, Amborella trichopoda* and *Marchantia polymorpha* (Fig S1 and Table S1 in SI Appendix). These four species represent critical evolutionary milestones across the history of land plants, ranging from the earliest terrestrial colonizers to highly specialized flowering plants.

To characterize structural organization across MADS-box proteins, we first reconstructed their phylogeny. The resulting tree was consistent with previous classifications and recovered the established clades, including SEP, AGL6, AP1/CAL, FYF, SOC1, AGL15, SVP, AP3, TT16, AG, PI, MIKC*, and FLC (Fig. 1) (19).

**Figure 1.**
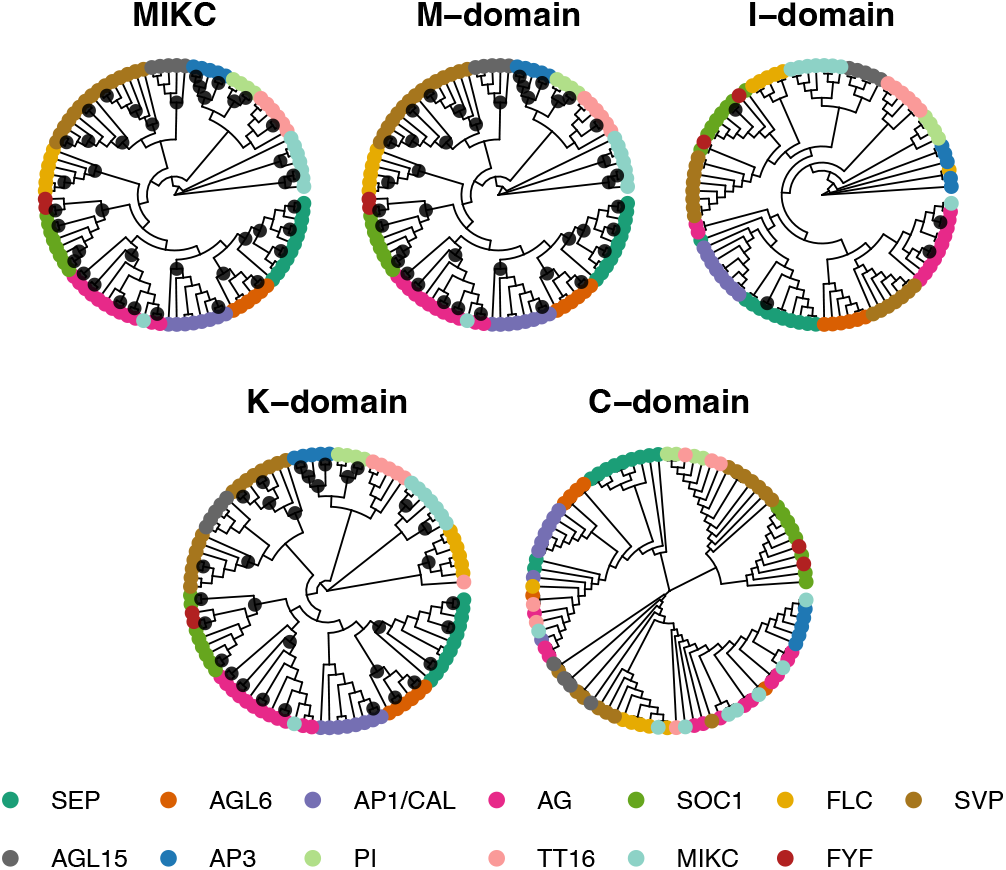
Phylogenetic reconstruction of MADS-box proteins and domain-specific trees. MADS-box proteins from *Arabidopsis thaliana, Marchantia polymorpha, Vitis vinifera*, and *Amborella trichopoda*, including MIKC* proteins (19, 31), were analyzed. Phylogenetic reconstruction based on aligned primary sequences using the JTT model and Neighbor-Joining method. Thirteen clades are resolved: SEP, AGL6/REDUCED BRANCHING, AP1/CAL, AG, SOC1, FLC, SVP, AGL15, AP3, PI, TT16, MIKC* and FYF. The MIKC* clade includes AtAGL30, AtAGL65, AtAGL67, AtAGL94, and AtAGL104. Topologies are shown for full-length proteins (MIKC) and for individual domains (M, I, K, and C). Black dots indicate nodes with bootstrap support ≥ 0.90.

We next examined predicted structural organization using AlphaFold-derived pLDDT scores, which provide residue-level confidence estimates (0–100). Residues with pLDDT ≥70 were classified as ordered, whereas those below 70 were considered disordered (20). Mapping pLDDT values along each protein revealed recurrent and clade-specific structural patterns (Fig. 2; Fig. S1 in SI Appendix).

**Figure 2.**
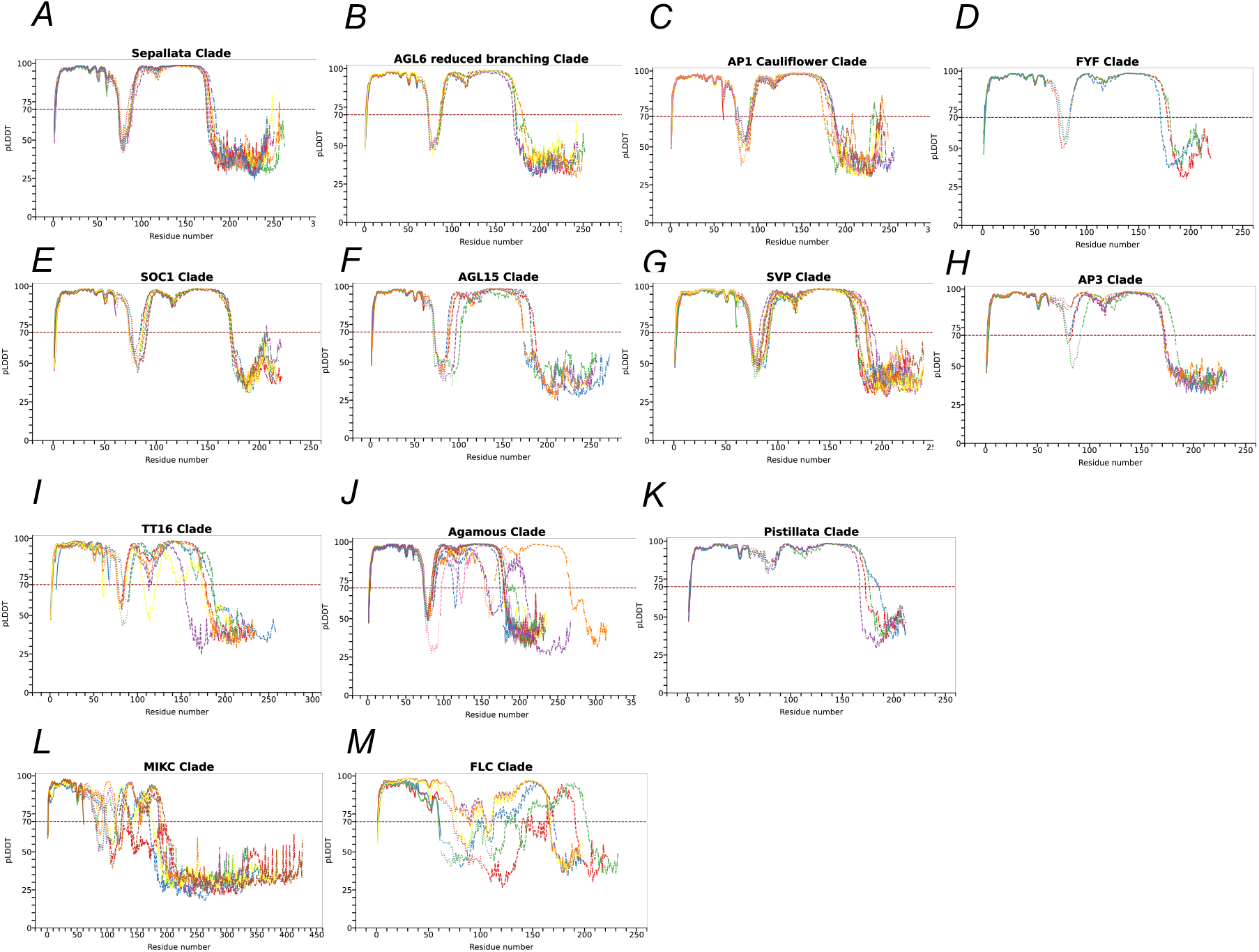
Domain-resolved pLDDT profiles reveal clade-specific structural patterns in MADS-box proteins. Per-residue predicted local distance difference test (pLDDT) scores derived from AlphaFold models are plotted against residue position. pLDDT values range from 0 to 100, with higher values indicating greater structural confidence. Residues with pLDDT ≥70 were classified as ordered, and residues with pLDDT <70 as disordered. The horizontal dashed line indicates the threshold (pLDDT = 70). Domain boundaries are indicated by line style: solid (M domain), dotted (I domain), dashed (K domain), and dash–dotted (C domain). (A–M) Representative clades: SEP (A), AGL6 (B), AP1/CAL (C), FYF (D), SOC1 (E), AGL15 (F), SVP (G), AP3 (H), TT16 (I), AG (J), PI (K), MIKC* (L), and FLC (M). Individual protein profiles for each clade are shown in Fig. S1.

Three broad structural classes emerged. The first group, comprising SEP, AGL6, AP1/CAL, FYF, SOC1, AGL15, and SVP (Fig. 2A–G), exhibited a conserved architectural profile. The MADS domain was strongly ordered, followed by a V-shaped disordered region spanning the I domain and the N-terminal portion of the K box. The central K domain was predominantly structured, whereas the C-terminal region was consistently disordered. Local reductions in structure (“wedges”) were reproducible across clades, particularly within the M domain and proximal K box, indicating conserved fine-scale modulation of structural stability.

A second group (AP3, TT16, and AG; Fig. 2H–J) displayed mixed structural features. Several AP3 orthologs lacked the canonical I-domain disorder, whereas Amborella paralogs retained it. In TT16 and AG clades, disordered segments within the K domain were expanded and, in some cases, extended to pLDDT values below 50. Notably, these structural differences occurred without substantial changes in domain length.

A third group (PI, MIKC*, and FLC; Fig. 2K–M) showed pronounced divergence. PI proteins were fully structured across M, I, and K domains. In contrast, MIKC* proteins showed fragmented structural organization, with alternating ordered and disordered segments following the M domain. The FLC clade displayed extensive disorder across the I and K domains. Together, these data reveal greater structural heterogeneity within the MADS-box family than previously appreciated.

We next examined whether apparent differences in I-domain structure reflected variation in domain length. Although some AP3 orthologs lacked disordered segments, comparison of M, I, and K domain lengths revealed minimal size differences among proteins. While the C domain varied considerably in length, it remained predominantly disordered across clades. Thus, ordered– disordered transitions are not explained by domain length but instead reflect intrinsic differences in structural organization.

### Phylogenetic signature is lost in the IDR I and C domains

To assess whether evolutionary signal is uniformly distributed across domains, we reconstructed phylogenies separately for the M, I, K, and C domains (Fig. 1).

The M domain recapitulated a phylogenetic reconstruction that was, at first sight, highly similar to that obtained using full-length MADS-box proteins (Robinson–Foulds distance = 0). Several clades remained intact, including AP3, PI, MIKC*, FLC, and AG. However, some M-domain sequences clustered into separate clades, such as AtGOA, AtXAL1, and VviXAL1, and others showed partial disruption, including SVP, AGL15, and AGL6. Although only a subset of nodes reached strong support (bootstrap ≥ 0.90), these corresponded to major clades, indicating that despite strong sequence conservation, the M domain retains a coherent phylogenetic signal sufficient to recover a largely stable topology.

Phylogenetic analysis of the I domain provided little support for previously defined clades, yet maintains some structural coherence. Although some proteins assigned to the same clade remained grouped, bootstrap support was generally low (<0.5).

In contrast, the K domain phylogeny was well resolved and showed strong statistical support. Its topology closely matched that obtained from complete coding sequences, as reflected by high Robinson–Foulds similarity, except for AtGOA, which clustered separately. Notably, the phylogenetic distances inferred from the MADS-box, M, and K domain trees were comparable.

Analysis of the C domain resulted in poor phylogenetic resolution. Clades lacked statistical support, indicating that although sequence divergence was sufficient to estimate genetic distances, it was insufficient to reconstruct a robust phylogeny.

These analyses indicate that phylogenetic signal is preserved in the structurally constrained M and K domains but largely diminished in the intrinsically disordered I and C domains. Evolutionary relationships inferred from full-length sequences therefore reflect uneven contributions from distinct structural regions.

### Constructing a metric for secondary structure-based distance

To quantify structural divergence, we developed a secondary structure distance (S2D) metric. AlphaFold models were retrieved in mmCIF format and converted to PDB files. Secondary structure was assigned using STRIDE (21). STRIDE provides the standard one-letter secondary structure code for every residue: (H) Alpha helix, (G) 3_10_ helix, (I) π-helix, (E) Extended conformation, (B/b) Isolated bridge, (T) Turn and (C) Coil. These assignments were filtered using pLDDT scores: residues with pLDDT <70 were classified as coil to avoid overinterpreting low-confidence predictions (Fig. 3).

**Figure 3.**
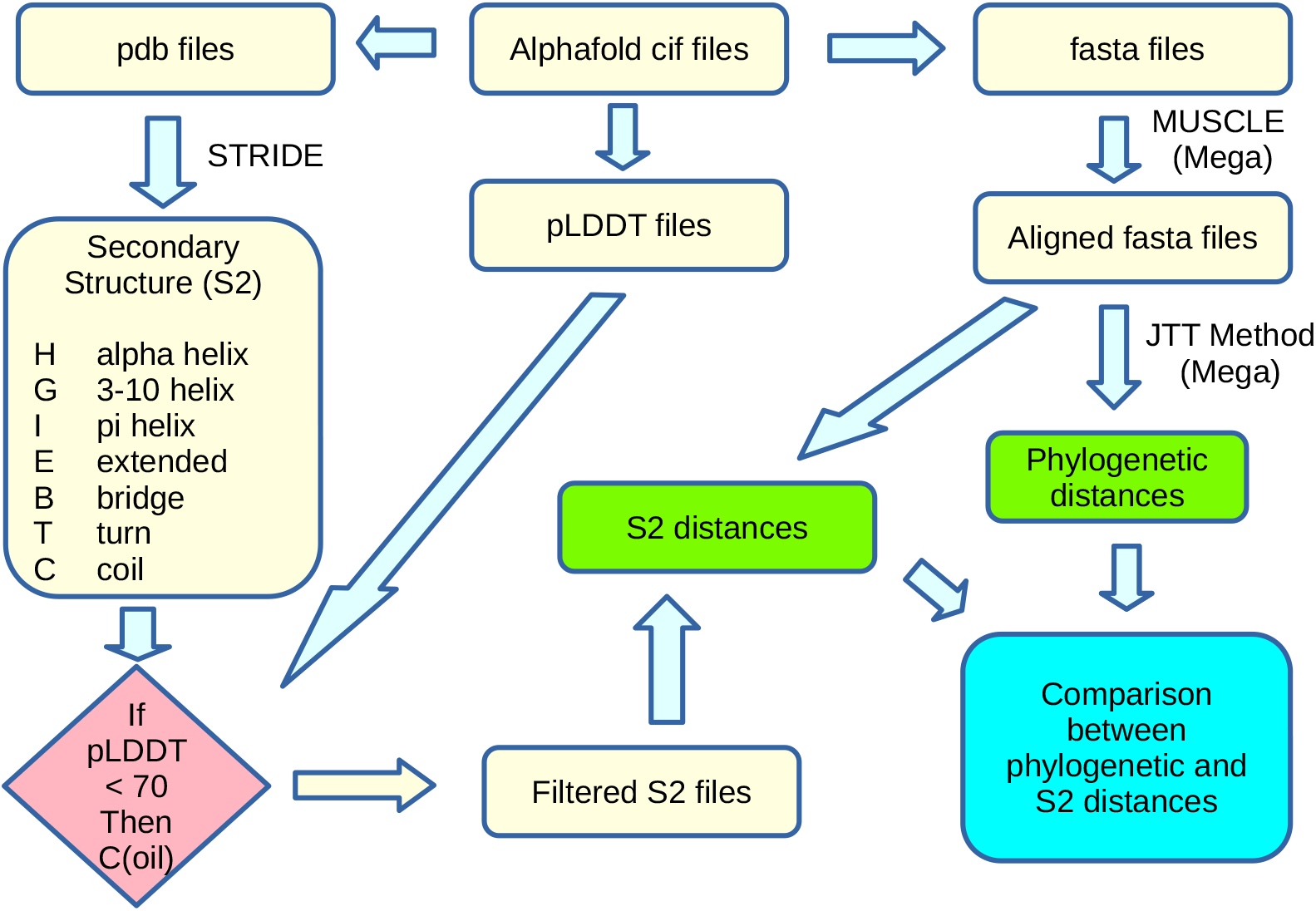
Workflow illustrating sequence alignment, secondary structure assignment, and calculation of phylogenetic and secondary structure distances (S2D).

Each aligned residue position was encoded by its STRIDE structural state. Application of the pLDDT filter increased coil assignments from 29.5% to 40.1%, indicating that approximately one-quarter of disordered regions were revealed only after confidence filtering. S2D between two proteins was calculated as the proportion of aligned residues differing in structural assignment, excluding gaps. Thus, S2D quantifies divergence in secondary structural organization independently of amino acid identity.

### Secondary structure distance reconstructions reveal domain-specific structural divergence

We produced a protein distance tree based on S2 distances using the complete MADS-box proteins. We found that the majority of the S2D clusters were combinations of proteins that appear separate in phylogenetic reconstructions (Fig 4). In contrast to sequence-based phylogeny, most S2D clusters comprised proteins that belong to separate phylogenetic clades. Only the MIKC* and FLC clades retained coherent clustering across both approaches. Notably, these two clades also exhibited large internal S2Ds and substantial divergence from the remainder of the family, underscoring their structural distinctiveness.

**Figure 4.**
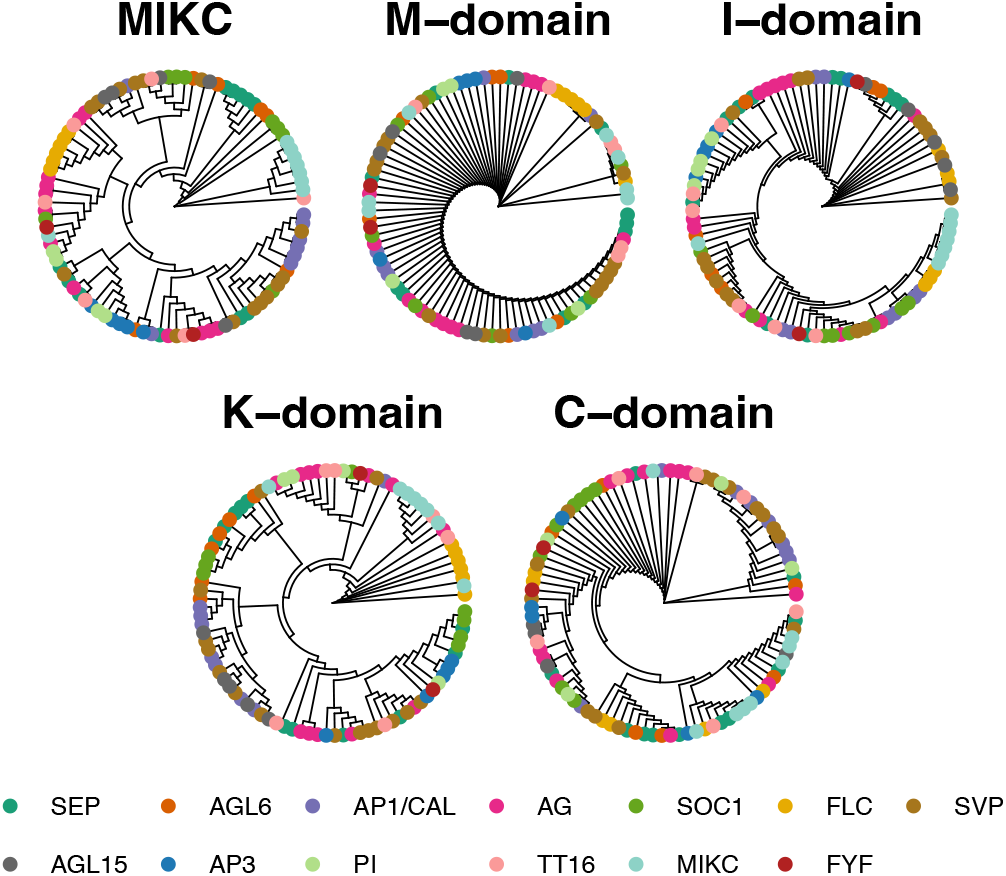
Trees based on secondary structure distances (S2D). Topologies are shown for full-length proteins (MIKC) and for individual domains (M, I, K, and C), enabling comparison of structural divergence across domains.

Domain-level S2D trees revealed distinct evolutionary trajectories. For the M domain, most proteins clustered tightly, consistent with low S2D values and strong structural conservation (Fig 4). However, several higher-distance clusters were detected, including a group containing all MIKC* proteins except AtMAF1 and mixed clusters combining members of TT16, MIKC*, SVP, and SEP clades. These results indicate that even minimal primary sequence variation within the highly conserved M domain can alter secondary structural organization.

The I-domain S2D tree showed pronounced fragmentation. The FLC clade separated into two distinct structural clusters: one comprising AtMAF4 and AtMAF5, and another containing AtFLC, AtMAF2, and AtMAF3, with substantial S2D between them (Fig 4). In contrast, the MIKC* clade remained structurally cohesive and positioned closer to the FLC cluster.

The K-domain S2D tree displayed greater structural dispersion than the M domain (Fig 4). Clusters frequently combined proteins from different phylogenetic clades, and S2D values were markedly higher than those observed for the M domain. Again, the largest structural divergences were observed within the MIKC* and FLC groups.

The C-domain S2D reconstruction revealed two major patterns (Fig. 4). Most proteins clustered with minimal structural divergence, whereas a subset formed distinct structural groups spanning multiple phylogenetic clades, including SVP, AtPI, AtGOA, AG, and AP1/CAL. Inspection of pLDDT profiles showed that several SVP proteins, AtPI, and AtGOA exhibit structured segments at the N-terminal region of the C domain. A similar feature was observed in VviAGL11, where residues 178– 195 showed elevated pLDDT values (68–88) (Fig. 2G–J). Conversely, AP1/CAL proteins displayed localized structural elements near the C-terminal end of the C domain.

Collectively, these analyses demonstrate that S2D reconstruction detects structural differentiation within individual domains, including subtle localized structural features, and reorganizes protein relationships independently of phylogenetic grouping. Structural divergence identified by S2D is not simply a function of global disorder but reflects domain-specific architectural variation.

### Phylogenetic and S2 distances show differing distribution patterns

We examined the pairwise distributions of phylogenetic distances for full-length proteins and for individual M, I, K, and C domains (n = 4,560 comparisons; Fig. S2A in SI Appendix). For complete MADS-box proteins, most pairwise distances fell between 1 and 2. As expected, the highly conserved M domain showed a narrower distribution, with most distances below 1. The K domain exhibited broader sequence divergence (0.5–3), indicating lower conservation relative to the M domain. In contrast, the intrinsically disordered I and C domains displayed markedly wider phylogenetic ranges (0.5–40 and 1.5–81, respectively), reflecting substantial sequence divergence.

We next examined the distribution of S2Ds (Fig. S2B in SI Appendix). For full-length proteins, most structural differences were below 15%, although a subset of comparisons reached ∼55%. The M domain again showed limited divergence, with most S2Ds below 5%, but a detectable tail extending to ∼20%, indicating that minor sequence variation can affect secondary structure.

The I and C domains displayed lower average S2Ds than full-length proteins but exhibited broad dispersion, with numerous comparisons showing substantial structural divergence. Notably, the K-domain S2D distribution closely resembled that of the complete proteins, suggesting that structural variation within the K box contributes disproportionately to overall secondary structure divergence. Together, these analyses indicate that sequence divergence and structural divergence follow distinct distribution patterns across domains, with the K domain emerging as a major driver of global structural variability.

### Comparisons between phylogeny and secondary structure distances identify outliers undergoing convergent and divergent evolution

To evaluate how structural divergence relates to sequence-based phylogeny, we plotted secondary structure distance (S2D) against phylogenetic distance for all 4,560 pairwise comparisons among MADS-box proteins (Fig. 5A). Rather than following a linear trend, the relationship was highly dispersed. Most protein pairs clustered at intermediate phylogenetic distances (1–1.5) with limited structural divergence (<15%). However, numerous comparisons deviated markedly from this pattern.

**Figure 5.**
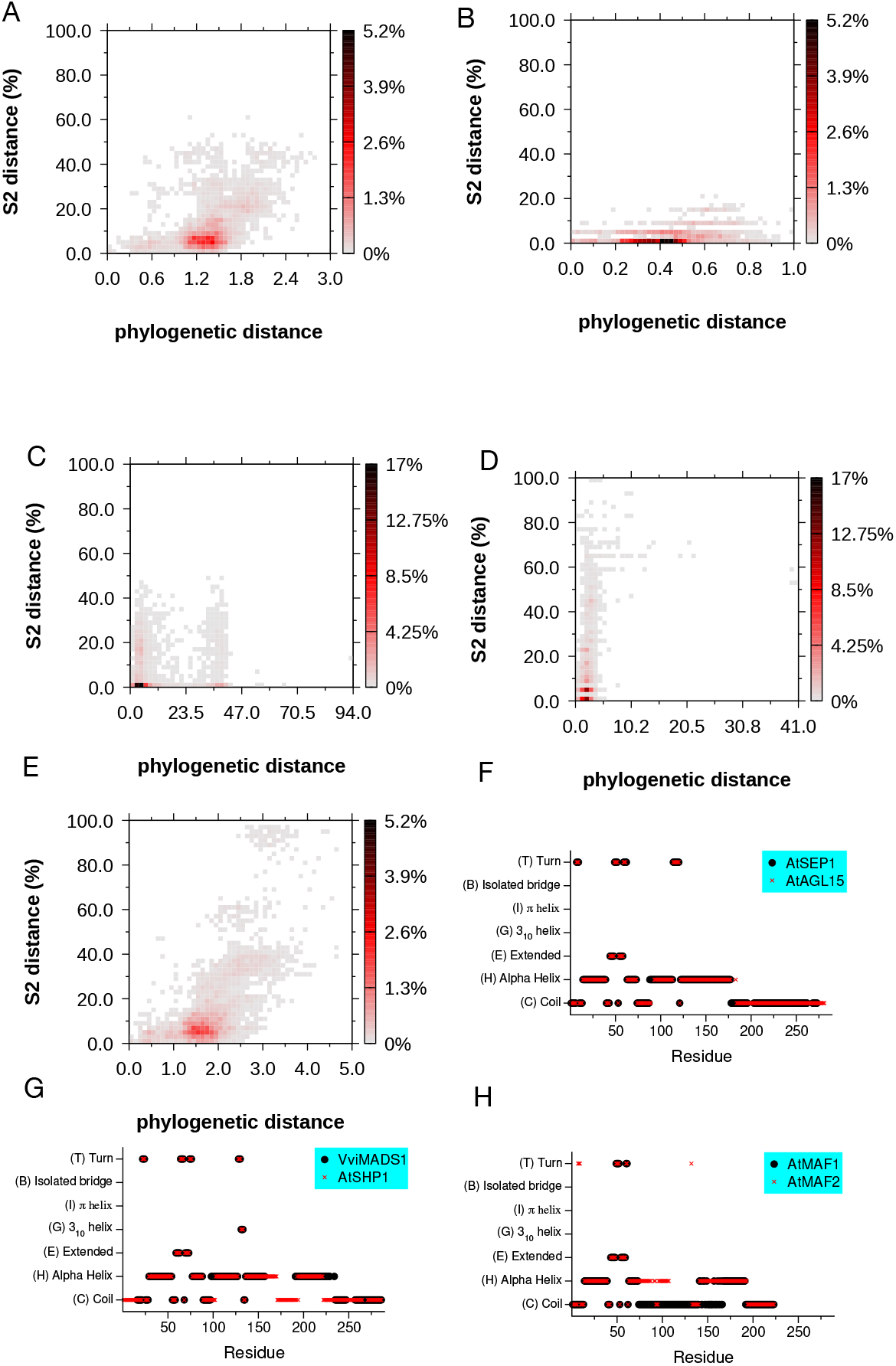
Integration of phylogenetic and secondary structure distances reveals convergent and divergent structural evolution. (A–E) Pairwise comparison of phylogenetic distance versus secondary structure distance (S2D) for all MADS-box protein pairs. Each point represents one pairwise comparison (n = 4560). Panels show complete proteins (A) and individual domains: M (B), I (C), K (D), and C (E). (F–H) Representative examples of structural outliers identified from pairwise comparisons. (F) STRIDE-derived secondary structure alignment of AtSEP1 and AtAGL15 illustrating low S2D despite substantial phylogenetic distance (convergent structural organization). (G) Comparison of AtMAF1 and AtMAF2 showing high S2D despite low phylogenetic distance (structural divergence). (H) Comparison of VviMADS1 and AtSHP1 showing structural divergence between closely related proteins. Secondary structure states are shown along aligned residue positions.

Strikingly, some closely related proteins (phylogenetic distance <1.0) exhibited substantial structural divergence (>40%), whereas other distantly related proteins (phylogenetic distance >2.0) showed minimal structural differences (<10%). These deviations indicate that secondary structure evolution is not uniformly coupled to primary sequence divergence.

Domain-level analyses revealed distinct evolutionary patterns. The M domain showed restricted variation, with phylogenetic distances generally below 1.2 and S2D values below 25% (Fig. 5B), consistent with strong structural conservation. In contrast, the I domain showed extreme dispersion, with phylogenetic distances extending up to 41 and S2D values approaching 100% (Fig. 5C). The K domain exhibited intermediate sequence divergence but substantial structural variability, with S2D values frequently exceeding 80% (Fig. 5D). The C domain displayed the broadest distribution, encompassing both low and high phylogenetic distances and structural divergences up to 80% (Fig. 5E). Together, these analyses indicate that individual domains contribute unevenly to overall structural divergence.

### Convergent structural evolution

We next examined extreme cases of structural convergence, defined as low S2D (<2%) despite substantial phylogenetic separation. A representative example within Arabidopsis is AtSEP1– AtAGL15 (phylogenetic distance 1.56; S2D 1.2%) (Fig 5F). STRIDE-derived motif comparison revealed nearly identical structural organization across M, I, and K domains, differing primarily in C-terminal length. Additional interspecific examples included AmAP3c–AtFUL, AmAP3c–VviSVP2, AtAGL15–VviAGL6, and VviMADS3–VviSVP2. These cases demonstrate that highly similar structural architectures can be maintained or re-established across distant phylogenetic lineages.

### Divergent structural evolution

Conversely, several closely related proteins displayed pronounced structural divergence. The paralogs VviSVPa–VviSVPb (phylogenetic distance 0.17) showed measurable structural differences (S2D 6.1%), and the interspecific pair VviMADS1–AtSHP1 (phylogenetic distance 0.31) exhibited S2D of 7.7% (Fig 5G), suggesting divergence not apparent from sequence comparison alone.

The most striking case involved the AtMAF paralogs. Although AtMAF1–AtMAF2 and AtMAF1– AtMAF3 are closely related (phylogenetic distances 0.36 and 0.32, respectively), both pairs exhibited substantial structural divergence (S2D 26.5%) (Fig 5H). In contrast, AtMAF2–AtMAF3 showed minimal structural difference (S2D 3.1%) despite comparable phylogenetic distance.

Structural motif comparison confirmed that AtMAF1 has undergone pronounced remodeling, particularly within the K domain, where loss of α-helical structure contrasts with the conserved helices in AtMAF2 and AtMAF3. These observations indicate lineage-specific structural innovation within an otherwise conserved clade.

Collectively, these results demonstrate that secondary structure evolution can proceed independently of primary sequence divergence, revealing both convergent and divergent evolutionary trajectories within the MADS-box family that are not evident from phylogeny alone.

### Identification of amino acid substitutions driving secondary structural change

Phylogenetic reconstruction assigns different weights to amino acid substitutions based on primary sequence conservation (2). To determine which substitutions most strongly influence secondary structure, we quantified the probability that a residue substitution results in a change of secondary structural state across aligned positions in the MADS-box family.

Substitutions involving tryptophan (W) showed the highest probability of inducing secondary structural changes (Fig. 6A). Additional substitutions with elevated structural impact included glutamic acid (E), cysteine (C), leucine (L), glycine (G), and histidine (H). In contrast, substitutions involving phenylalanine (F), valine (V), proline (P), and threonine (T) exhibited comparatively low probabilities of altering secondary structure.

**Figure 6.**
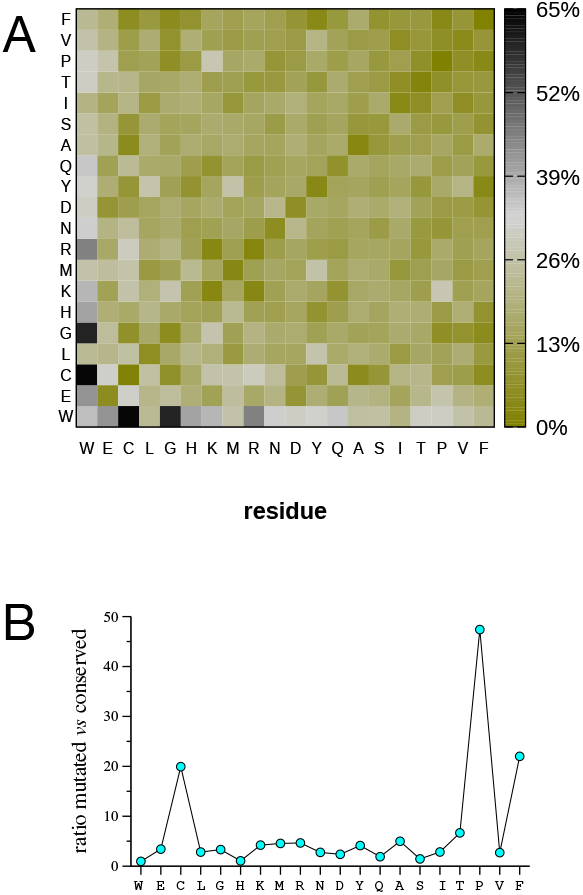
Amino acid substitutions exhibit differential effects on secondary structure. (A) Heatmap showing the percentage of secondary structure changes (S2D transitions) observed for each pairwise amino acid substitution across aligned positions. Each cell represents the proportion of cases in which substitution of one residue by another is associated with a change in STRIDE-assigned secondary structure state. (B) Ratio of mutation-induced structural change relative to intrinsic structural variability. For each amino acid, the probability that a substitution alters secondary structure is normalized by the variability observed for conserved residues. Higher values indicate substitutions with disproportionately strong impact on secondary structure.

Notably, cysteine, tryptophan, and histidine are among the most conserved residues in BLOSUM matrices, suggesting a convergence between primary sequence conservation and structural constraint. However, our analysis further highlights the structural sensitivity associated with glutamate and glycine substitutions, underscoring their contribution to secondary structural stability. We next examined conformational variability in the absence of substitution (diagonal comparisons) (Fig 6B). Tryptophan displayed the highest intrinsic structural variability (36%), whereas most other residues exhibited substantially lower variability, ranging from 13% for histidine to 0.3% for proline. Thus, conserved tryptophan residues can adopt multiple structural contexts, whereas conserved proline residues are typically associated with a stable structural state.

To disentangle intrinsic flexibility from mutation-driven effects, we calculated the ratio between the probability of structural change following substitution and the variability observed in conserved residues (Fig. 6B). This analysis revealed that mutations involving proline (47.4), phenylalanine (21.7), and cysteine (19.9) exert the strongest relative impact on secondary structure. In contrast, substitutions involving tryptophan (0.96), histidine (1.06), and serine (1.4) showed minimal relative effect.

Together, these results quantify the differential influence of individual amino acids on secondary structural stability and reveal that the structural consequences of mutation are not uniformly predicted by sequence conservation alone.

## Discussion

In this work, we developed a secondary structure–based distance metric (S2D) to quantify protein similarity beyond primary sequence comparisons. The rationale emerged from the observation that phylogenetic reconstructions of individual MADS-box domains yielded inconsistent results. While the conserved M and K domains produced robust clade separation, the I and C domains lost phylogenetic resolution and could not be unequivocally assigned to clades. This loss of phylogenetic signature reflects extensive primary sequence divergence. However, functional studies indicate that C domains are required for ternary complex formation (22, 23), suggesting that functional conservation may persist despite sequence divergence. These observations indicate that phylogenetic distance alone is not a reliable proxy for functional conservation in intrinsically disordered domains.

Our S2D metric was computed by comparing paired amino acids aligned at the primary sequence level and quantifying differences in their secondary structure states. This approach ensures that S2 comparisons remain deterministic and position-specific. Phylogenetic reconstructions correctly identify conserved folded domains, but the absence of phylogenetic signal in the I and C domains contrasts with their conserved classification as intrinsically disordered regions (24). This supports the view that IDRs are evolutionarily maintained due to functional constraints related to protein– protein interactions and structural flexibility, even when their primary sequences diverge extensively.

Comparison of phylogenetic distances across domains confirmed that M and K domains are more conserved at the primary sequence level than I and C domains. In contrast, S2D values showed a broader and more heterogeneous distribution across both full-length proteins and individual domains. Importantly, S2 similarity cannot be inferred from phylogenetic proximity. Single amino acid substitutions may have minimal impact on phylogenetic reconstruction yet substantially alter local secondary structure context.

The combined analysis of phylogenetic and S2 distances reveals cases of structural divergence and convergence. Protein pairs with low phylogenetic distance but high S2D likely represent divergent evolution at the secondary structural level. Conversely, pairs with high phylogenetic distance but low S2D suggest structural convergence. These discrepancies illustrate that evolutionary relationships inferred from primary sequence do not fully capture structural evolutionary trajectories.

At the residue scale, we further show that amino acid substitutions differ markedly in their probability of altering secondary structure context. Certain substitutions disproportionately reconfigure S2 states, whereas others preserve structural context despite sequence divergence. Residue-specific flexibility patterns indicate that some amino acids maintain stable structural assignments, while others occupy variable S2 states. These findings reveal structural constraints that are not captured by substitution matrices used in sequence-based phylogeny.

Taken together, our results demonstrate that protein evolution operates across at least two partially independent layers: primary sequence and secondary structure. Sequence divergence does not necessarily imply structural divergence, and structural similarity may persist despite loss of phylogenetic signal. This is particularly evident in intrinsically disordered regions, where tertiary-structure–based comparisons fail and primary sequence appears highly divergent. By ignoring secondary structure transitions, classical phylogenetic approaches overlook a substantial fraction of structural evolutionary dynamics.

We therefore propose that secondary structure represents a necessary evolutionary dimension. Integrating S2-based distances with primary sequence phylogeny enables detection of structural convergence and divergence that are invisible to sequence-based methods alone. Rather than replacing classical phylogeny, S2D expands the evolutionary framework by incorporating ordered and disordered structural states into comparative analysis. This integration provides a more complete representation of protein evolutionary history.

## Materials and Methods

### Structural data retrieval and secondary structure assignment

Crystallographic Information Files (mmCIFs) of the selected proteins were downloaded from the AlphaFold Protein Structure Database (AFDB)(25, 26). Protein Data Bank (PDB), FASTA and predicted local distance difference test (pLDDT) (11) files were generated from the mmCIFs using *cif2pdb* (27) and *cif2csv* implemented in Ubuntu 22.04.

Secondary structures for every residue was assigned with STRIDE (8) using the PDB files as input. STRIDE provides the standard one-letter secondary structure code: (H) Alpha helix, (G) 310 helix, (I) π-helix, (E) Extended conformation, (B/b) Isolated bridge, (T) Turn and (C) Coil. These assignments were subsequently filtered using the pLDDT scores. Assignments were filtered using pLDDT scores. Residues with pLDDT ≥70 were retained as predicted, whereas residues with pLDDT <70 were reassigned as coil (C), as pLDDT values above 70 generally indicate reliable backbone prediction (11).

### Sequence alignment and phylogenetic analysis

Protein sequences were aligned with MUSCLE (28) implemented in MEGA (v11.0.13)(29) using default parameters. Phylogenetic distances were evaluated using the Jones-Taylor-Thornton (JTT) substitution model also included in MEGA.

Distance between trees was assessed using the Robinson-Foulds algorithm using the APE R package (30).

### Secondary structure distance (S2D) calculation

Secondary structure distance (S2D) between two proteins was calculated from aligned sequences. For each pairwise comparison, aligned residue positions (excluding gaps) were evaluated, and the proportion of positions with differing secondary structure assignments was computed as:

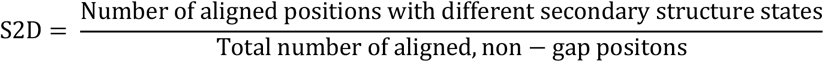

S2D values were expressed as percentages and treated as pairwise structural distances.

### Tree reconstruction

Phylogenetic distances and S2Ds were visualized using trees constructed with the Neighbour-joining (NJ) (5) statistical method also included in MEGA using 1000 bootstraps.

The complete bioinformatic flow is deposited in Github (https://github.com/abastidap/S2D), using the command line MEGA-CC.

## Acknowledgments

This work was partially supported by the Fundación Séneca– Agencia de Ciencia y Tecnología de la Región de Murcia (FSRM/10.13039/100007801) under Project 22570/PI/24.

