## Supplemental data for "Secondary structure distances reveal a new dimension of protein evolution"

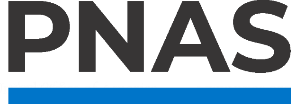

**Supporting Information for**

Secondary structure distances reveal a new dimension of protein evolution

Adolfo Bastida^a,1^, Ana María Muñoz-Morales^b^, and Marcos Egea-Cortines^c,1^

^a^Departamento de Quimica Física, Universidad de Murcia. 30100 Murcia, Spain. ^b^Plataforma de Big Data, IA, Bioestadística y Bioinformática, Instituto de Investigación Sanitaria La Fe (IIS La Fe), Valencia, Spain. ^c^Instituto de Biotecnología Vegetal, Universidad Politécnica de Cartagena member of European University of Technology, 30202 Cartagena, Spain

To whom correspondence may be addressed. Adolfo Bastida, Marcos Egea-Cortines.

**This PDF file includes:**

Figures S1 to S2

Tables S1

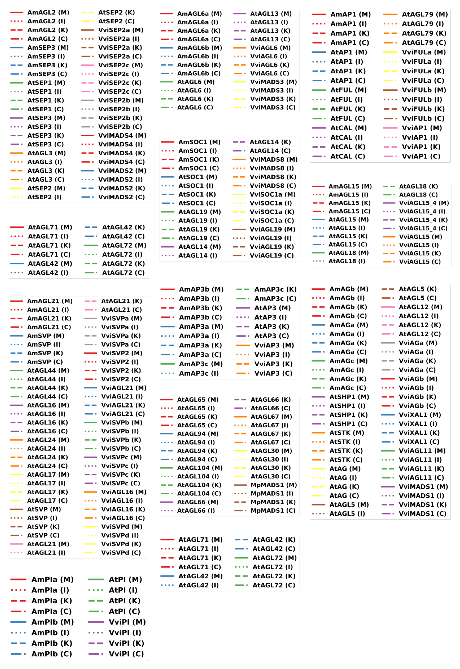

Fig. S1.Protein legends corresponding to Figure 2. The legend of the different clades correspond to (A) SEP clade (B) AGL6-reduced branching clade (C) AP1/CAL clade, (D) FYF clade, (E) SOC1 clade, (F) SVP clade (G) AP3 clade, (H) TT16 clade, (I) AGAMOUS clade, (J) PI clade (L) MIKC clade , (M) FLC clade.

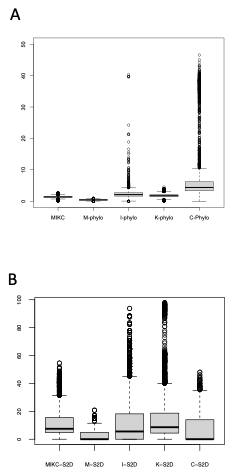

Fig. S2.Pairwise distribution of phylogenetic (A) and S2 distances (B) of the complete MADS box proteins, and the M, I, K and C domains. The data corresponds to 4560 comparisons using the dataset from table S1.

Tables

Table S1. Complete dataset with accession numbers

| **Clade** | **Organism** | **Protein name** | **iTAK, TAIR**  **or PlanTFDB Identifier** | **NCBI Identifier** | **AlphaFold Identifier** | **Identifier Uniprot** | **Gene name** |
| --- | --- | --- | --- | --- | --- | --- | --- |
| Sepallata | *Vitis vinifera* | Agamous-like MADS-box protein MADS2 | - | [XP_01908019](https://www.ncbi.nlm.nih.gov/protein/XP_019080193.1) [3.1](https://www.ncbi.nlm.nih.gov/protein/XP_019080193.1) | [VvSEP1](https://alphafold.ebi.ac.uk/search/text/VvSEP1) | [Q8LLR2](https://www.uniprot.org/uniprotkb/Q8LLR2/entry) | VviMADS2 |
| Sepallata | Vitis vinifera | Uncharacterized protein | - | [CAN84110.1](https://www.ncbi.nlm.nih.gov/protein/CAN84110.1) | [VITISV_0361](https://alphafold.ebi.ac.uk/search/text/VITISV_036170) [70](https://alphafold.ebi.ac.uk/search/text/VITISV_036170) | [A5C952](https://www.uniprot.org/uniprotkb/A5C952/entry) | VviSEP2a |
| Sepallata | *Vitis Vinifera* | Uncharacterized protein | [GSVIVT010](http://itak.feilab.net/cgi-bin/itak/db_gene_seq.cgi?trans_ID=GSVIVT01012249001) [12249001](http://itak.feilab.net/cgi-bin/itak/db_gene_seq.cgi?trans_ID=GSVIVT01012249001) | [XP_00226341](https://www.ncbi.nlm.nih.gov/protein/XP_002263410.2) [0.2](https://www.ncbi.nlm.nih.gov/protein/XP_002263410.2) | [VIT_01s0011](https://alphafold.ebi.ac.uk/search/text/VIT_01s0011g00110) [g00110](https://alphafold.ebi.ac.uk/search/text/VIT_01s0011g00110) | [D7T9Z7](https://www.uniprot.org/uniprotkb/D7T9Z7/entry) | VviSEP2b |
| Sepallata | *Vitis vinifera* | Uncharacterized protein | [GSVIVT010](http://itak.feilab.net/cgi-bin/itak/db_gene_seq.cgi?trans_ID=GSVIVT01008139001) [08139001](http://itak.feilab.net/cgi-bin/itak/db_gene_seq.cgi?trans_ID=GSVIVT01008139001) | [XP_00226303](https://www.ncbi.nlm.nih.gov/protein/XP_002263039.1) [9.1](https://www.ncbi.nlm.nih.gov/protein/XP_002263039.1) | [VIT_17s0000](https://alphafold.ebi.ac.uk/search/text/VIT_17s0000g05000) [g05000](https://alphafold.ebi.ac.uk/search/text/VIT_17s0000g05000) | [D7SIM7](https://www.uniprot.org/uniprotkb/D7SIM7/entry) | VviSEP2c |
| Sepallata | *Vitis vinifera* | Agamous-like MADS-box protein MADS4 | [GSVIVT010](http://itak.feilab.net/cgi-bin/itak/db_gene_seq.cgi?trans_ID=GSVIVT01010521001) [10521001](http://itak.feilab.net/cgi-bin/itak/db_gene_seq.cgi?trans_ID=GSVIVT01010521001) | [CBI27678.3](https://www.ncbi.nlm.nih.gov/protein/CBI27678.3) | [MADS4_VIT](https://alphafold.ebi.ac.uk/search/text/MADS4_VITVI) [VI](https://alphafold.ebi.ac.uk/search/text/MADS4_VITVI) | [Q8LLR0](https://www.uniprot.org/uniprotkb/Q8LLR0/entry) | VviMADS4 |
| Sepallata | *Arabidopsis thaliana* | Developmental  protein SEPALLATA 3 | [AT1G24260.](https://www.arabidopsis.org/servlets/TairObject?id=30252&type=locus)  [2](https://www.arabidopsis.org/servlets/TairObject?id=30252&type=locus) | [NP_564214.2](https://www.ncbi.nlm.nih.gov/protein/NP_564214.2) | [SEP3_ARAT](https://alphafold.ebi.ac.uk/search/text/SEP3_ARATH) [H](https://alphafold.ebi.ac.uk/search/text/SEP3_ARATH) | [O22456](https://www.uniprot.org/uniprotkb/O22456/entry) | AtSEP3 |
| Sepallata | *Amborella trichopoda* | Putative MADS-domain  transcription factor AGL9 | [evm_27.mod](http://itak.feilab.net/cgi-bin/itak/db_gene_seq.cgi?trans_ID=evm_27.model.AmTr_v1.0_scaffold00013.53) [el.AmTr_v1.](http://itak.feilab.net/cgi-bin/itak/db_gene_seq.cgi?trans_ID=evm_27.model.AmTr_v1.0_scaffold00013.53)  [0_scaffold00](http://itak.feilab.net/cgi-bin/itak/db_gene_seq.cgi?trans_ID=evm_27.model.AmTr_v1.0_scaffold00013.53) [013.53](http://itak.feilab.net/cgi-bin/itak/db_gene_seq.cgi?trans_ID=evm_27.model.AmTr_v1.0_scaffold00013.53) | [NP_00129276](https://www.ncbi.nlm.nih.gov/protein/NP_001292763.1) [3.1](https://www.ncbi.nlm.nih.gov/protein/NP_001292763.1) | [AMTR_s000](https://alphafold.ebi.ac.uk/search/text/AMTR_s00013p00103080) [13p0010308](https://alphafold.ebi.ac.uk/search/text/AMTR_s00013p00103080)  [0](https://alphafold.ebi.ac.uk/search/text/AMTR_s00013p00103080) | [W1PRJ9](https://www.uniprot.org/uniprotkb/W1PRJ9/entry) | AmSEP3 |
| Sepallata | *Amborella trichopoda* | AGL2 | [evm_27.mod](http://itak.feilab.net/cgi-bin/itak/db_gene_seq.cgi?trans_ID=evm_27.model.AmTr_v1.0_scaffold00047.121) [el.AmTr_v1.](http://itak.feilab.net/cgi-bin/itak/db_gene_seq.cgi?trans_ID=evm_27.model.AmTr_v1.0_scaffold00047.121) [0_scaffold00](http://itak.feilab.net/cgi-bin/itak/db_gene_seq.cgi?trans_ID=evm_27.model.AmTr_v1.0_scaffold00047.121) [047.121](http://itak.feilab.net/cgi-bin/itak/db_gene_seq.cgi?trans_ID=evm_27.model.AmTr_v1.0_scaffold00047.121) | [NP_00129275](https://www.ncbi.nlm.nih.gov/protein/NP_001292758.1) [8.1](https://www.ncbi.nlm.nih.gov/protein/NP_001292758.1) | [Q5D725_AM](https://alphafold.ebi.ac.uk/search/text/Q5D725_AMBTC) [BTC](https://alphafold.ebi.ac.uk/search/text/Q5D725_AMBTC) | [Q5D725](https://www.uniprot.org/uniprotkb/Q5D725/entry) | AmAGL2 |
| Sepallata | *Arabidopsis thaliana* | Developmental protein  SEPALLATA 2 | [AT3G02310.](https://www.arabidopsis.org/servlets/TairObject?id=35984&type=locus)  [1](https://www.arabidopsis.org/servlets/TairObject?id=35984&type=locus) | [NP_186880.1](https://www.ncbi.nlm.nih.gov/protein/NP_186880.1) | [SEP2_ARAT](https://alphafold.ebi.ac.uk/search/text/SEP2_ARATH) [H](https://alphafold.ebi.ac.uk/search/text/SEP2_ARATH) | [P29384](https://www.uniprot.org/uniprotkb/P29384/entry) | AtSEP2 |
| Sepallata | *Arabidopsis thaliana* | K-box region and MADS-box transcription  factor family protein | [AT5G15800.](https://www.arabidopsis.org/servlets/TairObject?id=130503&type=locus)  [2](https://www.arabidopsis.org/servlets/TairObject?id=130503&type=locus) | [NP_00111923](https://www.ncbi.nlm.nih.gov/protein/NP_001119230.1) [0.1](https://www.ncbi.nlm.nih.gov/protein/NP_001119230.1) | [F4KB90_AR](https://alphafold.ebi.ac.uk/search/text/F4KB90_ARATH) [ATH](https://alphafold.ebi.ac.uk/search/text/F4KB90_ARATH) | [F4KB90](https://www.uniprot.org/uniprotkb/F4KB90/entry) | AtSEP1 |
| Sepallata | *Arabidopsis thaliana* | Agamous-like  MADS-box protein AGL3 | [AT2G03710.](https://www.arabidopsis.org/servlets/TairObject?id=32207&type=locus)  [1](https://www.arabidopsis.org/servlets/TairObject?id=32207&type=locus) | [NP_178466.1](https://www.ncbi.nlm.nih.gov/protein/NP_178466.1) | [AGL3_ARAT](https://alphafold.ebi.ac.uk/search/text/AGL3_ARATH) [H](https://alphafold.ebi.ac.uk/search/text/AGL3_ARATH) | [P29383](https://www.uniprot.org/uniprotkb/P29383/entry) | AtAGL3 |
| AGL6  reduced branching | *Amborella trichopoda* | AGL6 | - | [AAY25580.1](https://www.ncbi.nlm.nih.gov/protein/AAY25580.1) | [Q2TDX2_AM](https://alphafold.ebi.ac.uk/search/text/Q2TDX2_AMBTC) [BTC](https://alphafold.ebi.ac.uk/search/text/Q2TDX2_AMBTC) | [Q2TDX2](https://www.uniprot.org/uniprotkb/Q2TDX2/entry) | AmAGL6a |
| AGL6  reduced branching | *Amborella trichopoda* | Uncharacterized protein | [evm_27.mod](http://itak.feilab.net/cgi-bin/itak/db_gene_seq.cgi?trans_ID=evm_27.model.AmTr_v1.0_scaffold00001.413) [el.AmTr_v1.](http://itak.feilab.net/cgi-bin/itak/db_gene_seq.cgi?trans_ID=evm_27.model.AmTr_v1.0_scaffold00001.413)  [0_scaffold00](http://itak.feilab.net/cgi-bin/itak/db_gene_seq.cgi?trans_ID=evm_27.model.AmTr_v1.0_scaffold00001.413) [001.413](http://itak.feilab.net/cgi-bin/itak/db_gene_seq.cgi?trans_ID=evm_27.model.AmTr_v1.0_scaffold00001.413) | [ERM96536.1](https://www.ncbi.nlm.nih.gov/protein/ERM96536.1) | [AMTR_s000](https://alphafold.ebi.ac.uk/search/text/AMTR_s00001p00267050) [01p0026705](https://alphafold.ebi.ac.uk/search/text/AMTR_s00001p00267050)  [0](https://alphafold.ebi.ac.uk/search/text/AMTR_s00001p00267050) | [W1NLY8](https://www.uniprot.org/uniprotkb/W1NLY8/entry) | AmAGL6b |
| AGL6  reduced branching | *Vitis vinifera* | Agamous-like MADS-box protein MADS3 | [GSVIVT010](http://itak.feilab.net/cgi-bin/itak/db_gene_seq.cgi?trans_ID=GSVIVT01027577001) [27577001](http://itak.feilab.net/cgi-bin/itak/db_gene_seq.cgi?trans_ID=GSVIVT01027577001) | [NP_00126811](https://www.ncbi.nlm.nih.gov/protein/NP_001268111.1) [1.1](https://www.ncbi.nlm.nih.gov/protein/NP_001268111.1) | [MADS3_VIT](https://alphafold.ebi.ac.uk/search/text/MADS3_VITVI) [VI](https://alphafold.ebi.ac.uk/search/text/MADS3_VITVI) | [Q8LLR1](https://www.uniprot.org/uniprotkb/Q8LLR1/entry) | VviMADS3 |
| AGL6  reduced branching | *Vitis vinifera* | Uncharacterized protein | [GSVIVT010](http://itak.feilab.net/cgi-bin/itak/db_gene_seq.cgi?trans_ID=GSVIVT01021303001) [213](http://itak.feilab.net/cgi-bin/itak/db_gene_seq.cgi?trans_ID=GSVIVT01021303001)  [03001](http://itak.feilab.net/cgi-bin/itak/db_gene_seq.cgi?trans_ID=GSVIVT01021303001) | [XP_05959583](https://www.ncbi.nlm.nih.gov/protein/XP_059595837.1?report=genbank&log%24=prottop&blast_rank=1&RID=UZAUAX1G013) [7.1](https://www.ncbi.nlm.nih.gov/protein/XP_059595837.1?report=genbank&log%24=prottop&blast_rank=1&RID=UZAUAX1G013) | [VIT_16s0022](https://alphafold.ebi.ac.uk/search/text/VIT_16s0022g02330)  [g](https://alphafold.ebi.ac.uk/search/text/VIT_16s0022g02330) [02330](https://alphafold.ebi.ac.uk/search/text/VIT_16s0022g02330) | [D7T1T8](https://www.uniprot.org/uniprotkb/D7T1T8/entry) | VviAGL6 |
| AGL6  reduced branching | *Arabidopsis thaliana* | Agamous-like MADS-box protein AGL13 | [AT3G61120.](https://www.arabidopsis.org/servlets/TairObject?id=39874&type=locus)  [1](https://www.arabidopsis.org/servlets/TairObject?id=39874&type=locus) | [NP_191671.1](https://www.ncbi.nlm.nih.gov/protein/NP_191671.1) | [AGL13_ARA](https://alphafold.ebi.ac.uk/search/text/AGL13_ARATH) [TH](https://alphafold.ebi.ac.uk/search/text/AGL13_ARATH) | [Q38837](https://www.uniprot.org/uniprotkb/Q38837/entry) | AtAGL13 |
| AGL6  reduced branching | *Arabidopsis thaliana* | Agamous-like MADS-box protein AGL6 | [AT2G45650.](https://www.arabidopsis.org/servlets/TairObject?id=32095&type=locus)  [1](https://www.arabidopsis.org/servlets/TairObject?id=32095&type=locus) | [NP_182089.1](https://www.ncbi.nlm.nih.gov/protein/NP_182089.1) | [AGL6_ARAT](https://alphafold.ebi.ac.uk/search/text/P29386) [H](https://alphafold.ebi.ac.uk/search/text/P29386) | [P29386](https://www.uniprot.org/uniprotkb/P29386/entry) | AtAGL6 |

| **Clade** | **Organism** | **Protein name** | **iTAK, TAIR**  **or PlanTFDB Identifier** | **NCBI**  **Identifier** | **AlphaFold Identifier** | **Identifier Uniprot** | **Gene name** |
| --- | --- | --- | --- | --- | --- | --- | --- |
| AP1  Cauliflower | *Arabidopsis thaliana* | Transcription factor CAULIFLOWER | [AT1G26310.](https://www.arabidopsis.org/servlets/TairObject?id=29732&type=locus)  [1](https://www.arabidopsis.org/servlets/TairObject?id=29732&type=locus) | [NP_564243.](https://www.ncbi.nlm.nih.gov/protein/NP_564243.1)  [1](https://www.ncbi.nlm.nih.gov/protein/NP_564243.1) | [CAL_ARAT](https://alphafold.ebi.ac.uk/search/text/CAL_ARATH) [H](https://alphafold.ebi.ac.uk/search/text/CAL_ARATH) | [Q39081](https://www.uniprot.org/uniprotkb/Q39081/entry) | AtCAL |
| AP1  Cauliflower | *Arabidopsis thaliana* | Floral homeotic protein APETALA 1 | [AT1G69120.](https://www.arabidopsis.org/servlets/TairObject?id=30417&type=locus)  [1](https://www.arabidopsis.org/servlets/TairObject?id=30417&type=locus) | [NP_177074.](https://www.ncbi.nlm.nih.gov/protein/NP_177074.1)  [1](https://www.ncbi.nlm.nih.gov/protein/NP_177074.1) | [AP1_ARATH](https://alphafold.ebi.ac.uk/search/text/AP1_ARATH) | [P35631](https://www.uniprot.org/uniprotkb/P35631/entry) | AtAP1 |
| AP1  Cauliflower | *Vitis vinifera* | Agamous-like MADS-box protein AP1 | [GSVIVT010](http://itak.feilab.net/cgi-bin/itak/db_gene_seq.cgi?trans_ID=GSVIVT01012250001) [1225](http://itak.feilab.net/cgi-bin/itak/db_gene_seq.cgi?trans_ID=GSVIVT01012250001)  [0001](http://itak.feilab.net/cgi-bin/itak/db_gene_seq.cgi?trans_ID=GSVIVT01012250001) | [AAT07447.1](https://www.ncbi.nlm.nih.gov/protein/AAT07447.1) | [AP1_VITVI](https://alphafold.ebi.ac.uk/search/text/AP1_VITVI) | [Q6E6S7](https://www.uniprot.org/uniprotkb/Q6E6S7/entry) | VviAP1 |
| AP1  Cauliflower | *Vitis vinifera* | Uncharacterized protein | [GSVIVT010](http://itak.feilab.net/cgi-bin/itak/db_gene_seq.cgi?trans_ID=GSVIVT01008140001) [0814](http://itak.feilab.net/cgi-bin/itak/db_gene_seq.cgi?trans_ID=GSVIVT01008140001)  [0001](http://itak.feilab.net/cgi-bin/itak/db_gene_seq.cgi?trans_ID=GSVIVT01008140001) | [XP_002263](https://www.ncbi.nlm.nih.gov/protein/XP_002263017.1) [017.1](https://www.ncbi.nlm.nih.gov/protein/XP_002263017.1) | [VIT_17s000](https://alphafold.ebi.ac.uk/search/text/VIT_17s0000g04990)  [0g](https://alphafold.ebi.ac.uk/search/text/VIT_17s0000g04990) [04990](https://alphafold.ebi.ac.uk/search/text/VIT_17s0000g04990) | [D7SIM8](https://www.uniprot.org/uniprotkb/D7SIM8/entry) | VviFULa |
| AP1  Cauliflower | *Vitis vinifera* | Agamous-like  MADS-box protein FUL-L | [GSVIVT010](http://itak.feilab.net/cgi-bin/itak/db_gene_seq.cgi?trans_ID=GSVIVT01036549001) [3654](http://itak.feilab.net/cgi-bin/itak/db_gene_seq.cgi?trans_ID=GSVIVT01036549001)  [9001](http://itak.feilab.net/cgi-bin/itak/db_gene_seq.cgi?trans_ID=GSVIVT01036549001) | [CBI16934.3](https://www.ncbi.nlm.nih.gov/protein/CBI16934.3) | [FULL_VITVI](https://alphafold.ebi.ac.uk/search/text/FULL_VITVI) | [D7SMN6](https://www.uniprot.org/uniprotkb/D7SMN6/entry) | VviFULb |
| AP1  Cauliflower | *Arabidopsis thaliana* | Agamous-like  MADS-box protein AGL8 | [AT5G60910.](https://www.arabidopsis.org/servlets/TairObject?id=134558&type=locus)  [1](https://www.arabidopsis.org/servlets/TairObject?id=134558&type=locus) | [NP_568929.](https://www.ncbi.nlm.nih.gov/protein/NP_568929.1)  [1](https://www.ncbi.nlm.nih.gov/protein/NP_568929.1) | [AGL8_ARAT](https://alphafold.ebi.ac.uk/search/text/AGL8_ARATH) [H](https://alphafold.ebi.ac.uk/search/text/AGL8_ARATH) | [Q38876](https://www.uniprot.org/uniprotkb/Q38876/entry) | AtAGL8 |
| AP1  Cauliflower | *Amborella trichopoda* | Uncharacterized protein | [evm_27.mod](http://itak.feilab.net/cgi-bin/itak/db_gene_seq.cgi?trans_ID=evm_27.model.AmTr_v1.0_scaffold00047.105) [el.AmTr_v1.](http://itak.feilab.net/cgi-bin/itak/db_gene_seq.cgi?trans_ID=evm_27.model.AmTr_v1.0_scaffold00047.105) [0_scaffold](http://itak.feilab.net/cgi-bin/itak/db_gene_seq.cgi?trans_ID=evm_27.model.AmTr_v1.0_scaffold00047.105)  [00047.105](http://itak.feilab.net/cgi-bin/itak/db_gene_seq.cgi?trans_ID=evm_27.model.AmTr_v1.0_scaffold00047.105) | [XP_0068563](https://www.ncbi.nlm.nih.gov/protein/XP_006856356.1) [56.1](https://www.ncbi.nlm.nih.gov/protein/XP_006856356.1) | [AMTR_s00](https://alphafold.ebi.ac.uk/search/text/AMTR_s00047p00181740) [047p001817](https://alphafold.ebi.ac.uk/search/text/AMTR_s00047p00181740)  [40](https://alphafold.ebi.ac.uk/search/text/AMTR_s00047p00181740) | [U5DBN6](https://www.uniprot.org/uniprotkb/U5DBN6/entry) | AmAP1 |
| AP1  Cauliflower | *Arabidopsis thaliana* | AGAMOUS-like 79 | [AT3G30260.](https://www.arabidopsis.org/servlets/TairObject?id=40572&type=locus)  [1](https://www.arabidopsis.org/servlets/TairObject?id=40572&type=locus) | [NP_189645.](https://www.ncbi.nlm.nih.gov/protein/NP_189645.2)  [2](https://www.ncbi.nlm.nih.gov/protein/NP_189645.2) | [Q7X9H6_AR](https://alphafold.ebi.ac.uk/search/text/Q7X9H6_ARATH) [ATH](https://alphafold.ebi.ac.uk/search/text/Q7X9H6_ARATH) | [Q7X9H6](https://www.uniprot.org/uniprotkb/Q7X9H6/entry) | AtAGL79 |
| Agamous | *Vitis vinifera* | Agamous | - | [NP_0012680](https://www.ncbi.nlm.nih.gov/protein/NP_001268097.1) [97.1](https://www.ncbi.nlm.nih.gov/protein/NP_001268097.1) | [D1MDP5_VI](https://alphafold.ebi.ac.uk/search/text/D1MDP5_VITVI) [TVI](https://alphafold.ebi.ac.uk/search/text/D1MDP5_VITVI) | [D1MDP5](https://www.uniprot.org/uniprotkb/D1MDP5/entry) | VviAGa |
| Agamous | *Vitis vinifera* | Uncharacterized protein | [GSVIVT010](http://itak.feilab.net/cgi-bin/itak/db_gene_seq.cgi?trans_ID=GSVIVT01021303001) [2130](http://itak.feilab.net/cgi-bin/itak/db_gene_seq.cgi?trans_ID=GSVIVT01021303001)  [3001](http://itak.feilab.net/cgi-bin/itak/db_gene_seq.cgi?trans_ID=GSVIVT01021303001) | [CBI30760.3](https://www.ncbi.nlm.nih.gov/protein/CBI30760.3) | [VIT_10s000](https://alphafold.ebi.ac.uk/search/text/VIT_10s0003g02070) [3g02070](https://alphafold.ebi.ac.uk/search/text/VIT_10s0003g02070) | [D7TJT8](https://www.uniprot.org/uniprotkb/D7TJT8/entry) | VviAGb |
| Agamous | *Vitis vinifera* | Agamous-like  MADS-box protein MADS1 | - | [NP_0012681](https://www.ncbi.nlm.nih.gov/protein/NP_001268105.1) [05.1](https://www.ncbi.nlm.nih.gov/protein/NP_001268105.1) | [MADS1_VIT](https://alphafold.ebi.ac.uk/search/text/MADS1_VITVI) [VI](https://alphafold.ebi.ac.uk/search/text/MADS1_VITVI) | [Q93XH4](https://www.uniprot.org/uniprotkb/Q93XH4/entry) | VviMADS1 |
| Agamous | *Arabidopsis thaliana* | Agamous-like MADS-box protein AGL5 | [AT2G42830.](https://www.arabidopsis.org/servlets/TairObject?id=33362&type=locus)  [1](https://www.arabidopsis.org/servlets/TairObject?id=33362&type=locus)  * | [NP_565986.](https://www.ncbi.nlm.nih.gov/protein/NP_565986.1)  [1](https://www.ncbi.nlm.nih.gov/protein/NP_565986.1) | [AGL5_ARAT](https://alphafold.ebi.ac.uk/search/text/AGL5_ARATH) [H](https://alphafold.ebi.ac.uk/search/text/AGL5_ARATH) | [P29385](https://www.uniprot.org/uniprotkb/P29385/entry) | AtAGL5 |
| Agamous | *Arabidopsis thaliana* | Floral homeotic protein AGAMOUS | [AT4G18960.](https://www.arabidopsis.org/servlets/TairObject?id=126907&type=locus)  [1](https://www.arabidopsis.org/servlets/TairObject?id=126907&type=locus)  * | [NP_567569.](https://www.ncbi.nlm.nih.gov/protein/NP_567569.3)  [3](https://www.ncbi.nlm.nih.gov/protein/NP_567569.3) | [AG_ARATH](https://alphafold.ebi.ac.uk/search/text/AG_ARATH) | [P17839](https://www.uniprot.org/uniprotkb/P17839/entry) | AtAG |
| Agamous | *Arabidopsis thaliana* | K-box region and MADS-box transcription factor family protein | [AT3G58780.](https://www.arabidopsis.org/servlets/TairObject?id=39912&type=locus)  [4](https://www.arabidopsis.org/servlets/TairObject?id=39912&type=locus) | [NP_0013257](https://www.ncbi.nlm.nih.gov/protein/NP_001325709.1) [09.1](https://www.ncbi.nlm.nih.gov/protein/NP_001325709.1) | [A0A1I9L](https://alphafold.ebi.ac.uk/search/text/A0A1I9LM26_ARATH) [M26_ARAT](https://alphafold.ebi.ac.uk/search/text/A0A1I9LM26_ARATH) [H](https://alphafold.ebi.ac.uk/search/text/A0A1I9LM26_ARATH) | [A0A1I9LM26](https://www.uniprot.org/uniprotkb/A0A1I9LM26/entry) | AtSHP1 |
| Agamous | *Amborella trichopoda* | Uncharacterized protein | [evm_27.mod](http://itak.feilab.net/cgi-bin/itak/db_gene_seq.cgi?trans_ID=evm_27.model.AmTr_v1.0_scaffold00071.203) [el.AmTr_v1.](http://itak.feilab.net/cgi-bin/itak/db_gene_seq.cgi?trans_ID=evm_27.model.AmTr_v1.0_scaffold00071.203) [0_scaffod](http://itak.feilab.net/cgi-bin/itak/db_gene_seq.cgi?trans_ID=evm_27.model.AmTr_v1.0_scaffold00071.203)  [00071.203](http://itak.feilab.net/cgi-bin/itak/db_gene_seq.cgi?trans_ID=evm_27.model.AmTr_v1.0_scaffold00071.203) | [XP_0068585](https://www.ncbi.nlm.nih.gov/protein/XP_006858589.1) [89.1](https://www.ncbi.nlm.nih.gov/protein/XP_006858589.1) | [AMTR_s000](https://alphafold.ebi.ac.uk/search/text/AMTR_s00071p00193200) [71p0019320](https://alphafold.ebi.ac.uk/search/text/AMTR_s00071p00193200)  [0](https://alphafold.ebi.ac.uk/search/text/AMTR_s00071p00193200) | [U5D3A6](https://www.uniprot.org/uniprotkb/U5D3A6/entry) | AmAGa |
| Agamous | *Amborella trichopoda* | AG | - | [AAY25577.1](https://www.ncbi.nlm.nih.gov/protein/AAY25577.1) | [Q2TDX5_A](https://alphafold.ebi.ac.uk/search/text/Q2TDX5_AMBTC) [MBTC](https://alphafold.ebi.ac.uk/search/text/Q2TDX5_AMBTC) | [Q2TDX5](https://www.uniprot.org/uniprotkb/Q2TDX5/entry) | AmAGb |
| Agamous | *Amborella trichopoda* | Uncharacterized protein | [evm_27.mod](http://itak.feilab.net/cgi-bin/itak/db_gene_seq.cgi?trans_ID=evm_27.model.AmTr_v1.0_scaffold00021.296) [el.AmTr_v1.](http://itak.feilab.net/cgi-bin/itak/db_gene_seq.cgi?trans_ID=evm_27.model.AmTr_v1.0_scaffold00021.296)  [0_scaffold00](http://itak.feilab.net/cgi-bin/itak/db_gene_seq.cgi?trans_ID=evm_27.model.AmTr_v1.0_scaffold00021.296) [021.296](http://itak.feilab.net/cgi-bin/itak/db_gene_seq.cgi?trans_ID=evm_27.model.AmTr_v1.0_scaffold00021.296) | [ERN14157.1](https://www.ncbi.nlm.nih.gov/protein/ERN14157.1) | [AMTR_s000](https://alphafold.ebi.ac.uk/search/text/AMTR_s00021p00254030) [21](https://alphafold.ebi.ac.uk/search/text/AMTR_s00021p00254030) [p00254030](https://alphafold.ebi.ac.uk/search/text/AMTR_s00021p00254030) | [W1PVX3](https://www.uniprot.org/uniprotkb/W1PVX3/entry) | AmAGc |

| **Clade** | **Organism** | **Protein name** | **iTAK, TAIR**  **or PlanTFDB Identifier** | **NCBI**  **Identifier** | **AlphaFold Identifier** | **Identifier Uniprot** | **Gene name** |
| --- | --- | --- | --- | --- | --- | --- | --- |
| Agamous | *Vitis vinifera* | Agamous-like MADS-box protein AGL11 | - | [XP_0106652](https://www.ncbi.nlm.nih.gov/protein/XP_010665297.1) [97.1](https://www.ncbi.nlm.nih.gov/protein/XP_010665297.1) | [AG11C_VIT](https://alphafold.ebi.ac.uk/search/text/AG11C_VITVI) [VI](https://alphafold.ebi.ac.uk/search/text/AG11C_VITVI) | [F6I457](https://www.uniprot.org/uniprotkb/F6I457/entry) | VviAGL11 |
| Agamous | *Arabidopsis thaliana* | K-box region and MADS-box transcription factor family protein | [AT4G09960.](https://www.arabidopsis.org/servlets/TairObject?id=130184&type=locus)  [4](https://www.arabidopsis.org/servlets/TairObject?id=130184&type=locus) | [NP_0013198](https://www.ncbi.nlm.nih.gov/protein/NP_001319889.1) [89.1](https://www.ncbi.nlm.nih.gov/protein/NP_001319889.1) | [F4JKV2_AR](https://alphafold.ebi.ac.uk/search/text/F4JKV2_ARATH) [ATH](https://alphafold.ebi.ac.uk/search/text/F4JKV2_ARATH) | [F4JKV2](https://www.uniprot.org/uniprotkb/F4JKV2/entry) | AtSTK |
| Agamous | *Arabidopsis thaliana* | Agamous-like MADS-box protein AGL12 | [AT1G71692.](https://www.arabidopsis.org/servlets/TairObject?id=226718&type=locus)  [1](https://www.arabidopsis.org/servlets/TairObject?id=226718&type=locus) | [NP_565022.](https://www.ncbi.nlm.nih.gov/protein/NP_565022.1)  [1](https://www.ncbi.nlm.nih.gov/protein/NP_565022.1) | [AGL12_ARA](https://alphafold.ebi.ac.uk/search/text/AGL12_ARATH) [TH](https://alphafold.ebi.ac.uk/search/text/AGL12_ARATH) | [Q38841](https://www.uniprot.org/uniprotkb/Q38841/entry) | AtXAL1 |
| Agamous | *Vitis vinifera* | Uncharacterized protein | [GSVIVT010](http://itak.feilab.net/cgi-bin/itak/db_gene_seq.cgi?trans_ID=GSVIVT01025916001) [2591](http://itak.feilab.net/cgi-bin/itak/db_gene_seq.cgi?trans_ID=GSVIVT01025916001)  [6001](http://itak.feilab.net/cgi-bin/itak/db_gene_seq.cgi?trans_ID=GSVIVT01025916001) | [XP_0022782](https://www.ncbi.nlm.nih.gov/protein/XP_002278239.1) [39.1](https://www.ncbi.nlm.nih.gov/protein/XP_002278239.1) | [VIT_18s004](https://alphafold.ebi.ac.uk/search/text/VIT_18s0041g02140)  [1g](https://alphafold.ebi.ac.uk/search/text/VIT_18s0041g02140) [02140](https://alphafold.ebi.ac.uk/search/text/VIT_18s0041g02140) | [D7U912](https://www.uniprot.org/uniprotkb/D7U912/entry) | VviXAL1 |
| SOC1 | *Vitis vinifera* | Suppressor of overexpression of CO 1 | - | [NP_0012679](https://www.ncbi.nlm.nih.gov/protein/NP_001267909.1) [09.1](https://www.ncbi.nlm.nih.gov/protein/NP_001267909.1) | [D1MDP7_VI](https://alphafold.ebi.ac.uk/search/text/D1MDP7_VITVI) [TVI](https://alphafold.ebi.ac.uk/search/text/D1MDP7_VITVI) | [D1MDP7](https://www.uniprot.org/uniprotkb/D1MDP7/entry) | VviSOC1a |
| SOC1 | *Vitis vinifera* | MADS-box protein | [GSVIVT010](http://itak.feilab.net/cgi-bin/itak/db_gene_seq.cgi?trans_ID=GSVIVT01027579001) [2757](http://itak.feilab.net/cgi-bin/itak/db_gene_seq.cgi?trans_ID=GSVIVT01027579001)  [9001](http://itak.feilab.net/cgi-bin/itak/db_gene_seq.cgi?trans_ID=GSVIVT01027579001) | [ABF56527.1](https://www.ncbi.nlm.nih.gov/protein/ABF56527.1) | [A1BQ41_VI](https://alphafold.ebi.ac.uk/search/text/A1BQ41_VITVI) [TVI](https://alphafold.ebi.ac.uk/search/text/A1BQ41_VITVI) | [A1BQ41](https://www.uniprot.org/uniprotkb/A1BQ41/entry) | VviMADS8 |
| SOC1 | *Arabidopsis thaliana* | MADS-box protein SOC1 | [AT2G45660.](https://www.arabidopsis.org/servlets/TairObject?id=26554&type=locus)  [1](https://www.arabidopsis.org/servlets/TairObject?id=26554&type=locus) | [NP_182090.](https://www.ncbi.nlm.nih.gov/protein/NP_182090.1)  [1](https://www.ncbi.nlm.nih.gov/protein/NP_182090.1) | [SOC1_ARA](https://alphafold.ebi.ac.uk/search/text/SOC1_ARATH) [TH](https://alphafold.ebi.ac.uk/search/text/SOC1_ARATH) | [O64645](https://www.uniprot.org/uniprotkb/O64645/entry) | AtSOC1 |
| SOC1 | *Amborella trichopoda* | Uncharacterized protein | [evm_27.](http://itak.feilab.net/cgi-bin/itak/db_gene_seq.cgi?trans_ID=evm_27.model.AmTr_v1.0_scaffold00001.409) [model.](http://itak.feilab.net/cgi-bin/itak/db_gene_seq.cgi?trans_ID=evm_27.model.AmTr_v1.0_scaffold00001.409)  [AmTr_v1.0_](http://itak.feilab.net/cgi-bin/itak/db_gene_seq.cgi?trans_ID=evm_27.model.AmTr_v1.0_scaffold00001.409)  [scaffold](http://itak.feilab.net/cgi-bin/itak/db_gene_seq.cgi?trans_ID=evm_27.model.AmTr_v1.0_scaffold00001.409) [00001.409](http://itak.feilab.net/cgi-bin/itak/db_gene_seq.cgi?trans_ID=evm_27.model.AmTr_v1.0_scaffold00001.409) | [XP_0068291](https://www.ncbi.nlm.nih.gov/protein/XP_006829116.1) [16.1](https://www.ncbi.nlm.nih.gov/protein/XP_006829116.1) | [AMTR_s000](https://alphafold.ebi.ac.uk/search/text/AMTR_s00001p00266470) [0](https://alphafold.ebi.ac.uk/search/text/AMTR_s00001p00266470)  [1p00266470](https://alphafold.ebi.ac.uk/search/text/AMTR_s00001p00266470) | [W1NL13](https://www.uniprot.org/uniprotkb/W1NL13/entry) | AmSOC1 |
| SOC1 | *Vitis vinifera* | Uncharacterized protein | - | [XP_0022756](https://www.ncbi.nlm.nih.gov/protein/XP_002275695.2) [95.2](https://www.ncbi.nlm.nih.gov/protein/XP_002275695.2) | [VIT_02s002](https://alphafold.ebi.ac.uk/search/text/VIT_02s0025g04650) [5](https://alphafold.ebi.ac.uk/search/text/VIT_02s0025g04650)  [g04650](https://alphafold.ebi.ac.uk/search/text/VIT_02s0025g04650) | [F6HUJ0](https://www.uniprot.org/uniprotkb/F6HUJ0/entry) | VviAGL19 |
| SOC1 | *Arabidopsis thaliana* | Agamous-like MADS-box protein AGL14 | [AT4G11880.](https://www.arabidopsis.org/servlets/TairObject?id=129748&type=locus)  [1](https://www.arabidopsis.org/servlets/TairObject?id=129748&type=locus) | [NP_0013199](https://www.ncbi.nlm.nih.gov/protein/NP_001319907.1) [07.1](https://www.ncbi.nlm.nih.gov/protein/NP_001319907.1) | [AGL14_ARA](https://alphafold.ebi.ac.uk/search/text/AGL14_ARATH) [TH](https://alphafold.ebi.ac.uk/search/text/AGL14_ARATH) | [Q38838](https://www.uniprot.org/uniprotkb/Q38838/entry) | AtAGL14 |
| SOC1 | *Arabidopsis thaliana* | Agamous-like MADS-box protein AGL19 | [AT4G22950.](https://www.arabidopsis.org/servlets/TairObject?id=128336&type=locus)  [1](https://www.arabidopsis.org/servlets/TairObject?id=128336&type=locus) | [NP_0013200](https://www.ncbi.nlm.nih.gov/protein/NP_001320037.1) [37.1](https://www.ncbi.nlm.nih.gov/protein/NP_001320037.1) | [AGL19_ARA](https://alphafold.ebi.ac.uk/search/text/AGL19_ARATH) [TH](https://alphafold.ebi.ac.uk/search/text/AGL19_ARATH) | [O82743](https://www.uniprot.org/uniprotkb/O82743/entry) | AtAGL19 |
| FYF | *Arabidopsis thaliana* | MADS-box protein AGL72 | [AT5G51860.](https://www.arabidopsis.org/servlets/TairObject?id=133487&type=locus)  [1](https://www.arabidopsis.org/servlets/TairObject?id=133487&type=locus) | [NP_199999.](https://www.ncbi.nlm.nih.gov/protein/NP_199999.1)  [1](https://www.ncbi.nlm.nih.gov/protein/NP_199999.1) | [AGL72_ARA](https://alphafold.ebi.ac.uk/search/text/AGL72_ARATH) [TH](https://alphafold.ebi.ac.uk/search/text/AGL72_ARATH) | [Q9FLH5](https://www.uniprot.org/uniprotkb/Q9FLH5/entry) | AtAGL72 |
| FYF | *Arabidopsis thaliana* | AGAMOUS-like 71 | [AT5G51870.](https://www.arabidopsis.org/servlets/TairObject?id=133625&type=locus)  [3](https://www.arabidopsis.org/servlets/TairObject?id=133625&type=locus) | [NP_0011905](https://www.ncbi.nlm.nih.gov/protein/NP_001190517) [17](https://www.ncbi.nlm.nih.gov/protein/NP_001190517).2 | [F4KEP6_AR](https://alphafold.ebi.ac.uk/search/text/F4KEP6_ARATH) [ATH](https://alphafold.ebi.ac.uk/search/text/F4KEP6_ARATH) | [F4KEP6](https://www.uniprot.org/uniprotkb/F4KEP6/entry) | AtAGL71 |

| **Clade** | **Organism** | **Protein name** | **iTAK, TAIR**  **or PlanTFDB Identifier** | **NCBI**  **Identifier** | **AlphaFold Identifier** | **Identifier Uniprot** | **Gene name** |
| --- | --- | --- | --- | --- | --- | --- | --- |
| FYF | *Arabidopsis thaliana* | MADS-box protein AGL42 | [AT5G62165.](https://www.arabidopsis.org/servlets/TairObject?id=500232022&type=locus)  [1](https://www.arabidopsis.org/servlets/TairObject?id=500232022&type=locus) | [NP_0010321](https://www.ncbi.nlm.nih.gov/protein/NP_001032123.1) [23.1](https://www.ncbi.nlm.nih.gov/protein/NP_001032123.1) | [AGL42_ARA](https://alphafold.ebi.ac.uk/search/text/AGL42_ARATH) [TH](https://alphafold.ebi.ac.uk/search/text/AGL42_ARATH) | [Q9FIS1](https://www.uniprot.org/uniprotkb/Q9FIS1/entry) | AtAGL42 |
| SOC1 | *Vitis vinifera* | Uncharacterized protein | [GSVIVT010](http://itak.feilab.net/cgi-bin/itak/db_gene_seq.cgi?trans_ID=GSVIVT01008560001) [0856](http://itak.feilab.net/cgi-bin/itak/db_gene_seq.cgi?trans_ID=GSVIVT01008560001)  [0001](http://itak.feilab.net/cgi-bin/itak/db_gene_seq.cgi?trans_ID=GSVIVT01008560001) | [CBI15681.3](https://www.ncbi.nlm.nih.gov/protein/CBI15681.3) | [VIT_17s000](https://alphafold.ebi.ac.uk/search/text/VIT_17s0000g01230) [0g01230](https://alphafold.ebi.ac.uk/search/text/VIT_17s0000g01230) | [D7SH01](https://www.uniprot.org/uniprotkb/D7SH01/entry) | VviSOC1b |
| AGL15 | *Vitis vinifera* | Agamous-like MADS-box protein AGL15 | - | [XP_0022785](https://www.ncbi.nlm.nih.gov/protein/XP_002278584.1) [84.1](https://www.ncbi.nlm.nih.gov/protein/XP_002278584.1) | [A0A438JK](https://alphafold.ebi.ac.uk/search/text/A0A438JKY7_VITVI) [Y7_VITVI](https://alphafold.ebi.ac.uk/search/text/A0A438JKY7_VITVI) | [A0A438JKY](https://www.uniprot.org/uniprotkb/A0A438JKY7/entry) [7](https://www.uniprot.org/uniprotkb/A0A438JKY7/entry) | VviAGL15_4 |
| AGL15 | *Vitis vinifera* | AGL15 | [GSVIVT010](http://itak.feilab.net/cgi-bin/itak/db_gene_seq.cgi?trans_ID=GSVIVT01001437001) [01437001](http://itak.feilab.net/cgi-bin/itak/db_gene_seq.cgi?trans_ID=GSVIVT01001437001) | [CBI28594.3](https://www.ncbi.nlm.nih.gov/protein/CBI28594.3) | [D7TDM1_VI](https://alphafold.ebi.ac.uk/search/text/D7TDM1_VITVI) [TVI](https://alphafold.ebi.ac.uk/search/text/D7TDM1_VITVI) | [D7TDM1](https://www.uniprot.org/uniprotkb/D7TDM1/entry) | VviAGL15 |
| AGL15 | *Arabidopsis thaliana* | AGAMOUS-like 15 | [AT5G13790.](https://www.arabidopsis.org/servlets/TairObject?id=135070&type=locus)  [2](https://www.arabidopsis.org/servlets/TairObject?id=135070&type=locus) | [NP_0013302](https://www.ncbi.nlm.nih.gov/protein/NP_001330207.1) [07.1](https://www.ncbi.nlm.nih.gov/protein/NP_001330207.1) | [A0A178UBU](https://alphafold.ebi.ac.uk/search/text/A0A178UBU5_ARATH) [5_ARATH](https://alphafold.ebi.ac.uk/search/text/A0A178UBU5_ARATH) | [A0A178UBU](https://www.uniprot.org/uniprotkb/A0A178UBU5/entry) [5](https://www.uniprot.org/uniprotkb/A0A178UBU5/entry) | AtAGL15 |
| AGL15 | *Amborella trichopoda* | Uncharacterized protein | [evm_27.mod](http://itak.feilab.net/cgi-bin/itak/db_gene_seq.cgi?trans_ID=evm_27.model.AmTr_v1.0_scaffold00053.185) [el.AmTr_v1.](http://itak.feilab.net/cgi-bin/itak/db_gene_seq.cgi?trans_ID=evm_27.model.AmTr_v1.0_scaffold00053.185) [0_scaffold00](http://itak.feilab.net/cgi-bin/itak/db_gene_seq.cgi?trans_ID=evm_27.model.AmTr_v1.0_scaffold00053.185)  [053.185](http://itak.feilab.net/cgi-bin/itak/db_gene_seq.cgi?trans_ID=evm_27.model.AmTr_v1.0_scaffold00053.185) | [ERN05188.1](https://www.ncbi.nlm.nih.gov/protein/ERN05188.1) | [AMTR_s000](https://alphafold.ebi.ac.uk/search/text/AMTR_s00053p00228660) [53p0022866](https://alphafold.ebi.ac.uk/search/text/AMTR_s00053p00228660)  [0](https://alphafold.ebi.ac.uk/search/text/AMTR_s00053p00228660) | [W1PB86](https://www.uniprot.org/uniprotkb/W1PB86/entry) | AmAGL15 |
| AGL15 | *Arabidopsis thaliana* | Agamous-like MADS-box protein AGL18 | [AT3G57390.](https://www.arabidopsis.org/servlets/TairObject?id=37043&type=locus)  [1](https://www.arabidopsis.org/servlets/TairObject?id=37043&type=locus) | [NP_191298.](https://www.ncbi.nlm.nih.gov/protein/NP_191298.1)  [1](https://www.ncbi.nlm.nih.gov/protein/NP_191298.1) | [AGL18_ARA](https://alphafold.ebi.ac.uk/search/text/AGL18_ARATH) [TH](https://alphafold.ebi.ac.uk/search/text/AGL18_ARATH) | [Q9M2K8](https://www.uniprot.org/uniprotkb/Q9M2K8/entry) | AtAGL18 |
| SVP | *Vitis vinifera* | Agamous-like MADS-box protein 24 | [GSVIVT010](http://itak.feilab.net/cgi-bin/itak/db_gene_seq.cgi?trans_ID=GSVIVT01001701001) [01701001](http://itak.feilab.net/cgi-bin/itak/db_gene_seq.cgi?trans_ID=GSVIVT01001701001) | [CBI35558.3](https://www.ncbi.nlm.nih.gov/protein/CBI35558.3) | [A0A3G2BC0](https://alphafold.ebi.ac.uk/search/text/A0A3G2BC07_VITVI) [7_VITVI](https://alphafold.ebi.ac.uk/search/text/A0A3G2BC07_VITVI) | [A0A3G2BC0](https://www.uniprot.org/uniprotkb/A0A3G2BC07/entry) [7](https://www.uniprot.org/uniprotkb/A0A3G2BC07/entry) | VviSVPa |
| SVP | *Vitis vinifera* | DAM3 | [GSVIVT010](http://itak.feilab.net/cgi-bin/itak/db_gene_seq.cgi?trans_ID=GSVIVT01011300001) [11300001](http://itak.feilab.net/cgi-bin/itak/db_gene_seq.cgi?trans_ID=GSVIVT01011300001) | [QBC35958.1](https://www.ncbi.nlm.nih.gov/protein/QBC35958.1/) | [VIT_15s010](https://alphafold.ebi.ac.uk/search/text/VIT_15s0107g00120) [7g00120](https://alphafold.ebi.ac.uk/search/text/VIT_15s0107g00120) | [D7U627](https://www.uniprot.org/uniprotkb/D7U627/entry) | VviSVPb |
| SVP | *Vitis vinifera* | Uncharacterized protein | [GSVIVT010](http://itak.feilab.net/cgi-bin/itak/db_gene_seq.cgi?trans_ID=GSVIVT01005934001) [05934001](http://itak.feilab.net/cgi-bin/itak/db_gene_seq.cgi?trans_ID=GSVIVT01005934001) | [XP_0190738](https://www.ncbi.nlm.nih.gov/protein/XP_019073897.1) [97.1](https://www.ncbi.nlm.nih.gov/protein/XP_019073897.1) | [VIT_00s031](https://alphafold.ebi.ac.uk/search/text/VIT_00s0313g00070) [3g00070](https://alphafold.ebi.ac.uk/search/text/VIT_00s0313g00070) | [D7U6A4](https://www.uniprot.org/uniprotkb/D7U6A4/entry) | VviSVPc |
| SVP | *Vitis vinifera* | SVP-like MADS-box protein | - | [AFC96914.1](https://www.ncbi.nlm.nih.gov/protein/AFC96914.1) | [H9CTT8_VI](https://alphafold.ebi.ac.uk/search/text/H9CTT8_VITVI) [TVI](https://alphafold.ebi.ac.uk/search/text/H9CTT8_VITVI) | [H9CTT8](https://www.uniprot.org/uniprotkb/H9CTT8/entry) | VviSVPd |
| SVP | *Vitis vinifera* | Agamous-like  MADS-box protein SVP | [GSVIVT010](http://itak.feilab.net/cgi-bin/itak/db_gene_seq.cgi?trans_ID=GSVIVT01009171001) [09171001](http://itak.feilab.net/cgi-bin/itak/db_gene_seq.cgi?trans_ID=GSVIVT01009171001) | [XP_0022856](https://www.ncbi.nlm.nih.gov/protein/XP_002285687.1) [87.1](https://www.ncbi.nlm.nih.gov/protein/XP_002285687.1) | [A0A3G2BC1](https://alphafold.ebi.ac.uk/search/text/A0A3G2BC13_VITVI) [3_VITVI](https://alphafold.ebi.ac.uk/search/text/A0A3G2BC13_VITVI) | [A0A3G2BC1](https://www.uniprot.org/uniprotkb/A0A3G2BC13/entry) [3](https://www.uniprot.org/uniprotkb/A0A3G2BC13/entry) | VviSVP2 |
| SVP | Arabidopsis thaliana | MADS-box protein SVP | [AT2G22540.](https://www.arabidopsis.org/servlets/TairObject?id=31694&type=locus)  [1](https://www.arabidopsis.org/servlets/TairObject?id=31694&type=locus) | [NP_0013245](https://www.ncbi.nlm.nih.gov/protein/NP_001324584.1) [84.1](https://www.ncbi.nlm.nih.gov/protein/NP_001324584.1) | [SVP_ARAT](https://alphafold.ebi.ac.uk/search/text/SVP_ARATH) [H](https://alphafold.ebi.ac.uk/search/text/SVP_ARATH) | [Q9FVC1](https://www.uniprot.org/uniprotkb/Q9FVC1/entry) | AtSVP |
| SVP | Arabidopsis thaliana | MADS-box protein AGL24 | [AT4G24540.](https://www.arabidopsis.org/servlets/TairObject?id=127586&type=locus)  [1](https://www.arabidopsis.org/servlets/TairObject?id=127586&type=locus) | [NP_194185.](https://www.ncbi.nlm.nih.gov/protein/NP_194185.1)  [1](https://www.ncbi.nlm.nih.gov/protein/NP_194185.1) | [AGL24_ARA](https://alphafold.ebi.ac.uk/search/text/AGL24_ARATH) [TH](https://alphafold.ebi.ac.uk/search/text/AGL24_ARATH) | [O82794](https://www.uniprot.org/uniprotkb/O82794/entry) | AtAGL24 |
| SVP | *Amborella trichopoda* | Uncharacterized protein | [evm_27.mod](http://itak.feilab.net/cgi-bin/itak/db_gene_seq.cgi?trans_ID=evm_27.model.AmTr_v1.0_scaffold00127.17) [el.AmTr_v1.](http://itak.feilab.net/cgi-bin/itak/db_gene_seq.cgi?trans_ID=evm_27.model.AmTr_v1.0_scaffold00127.17) [0_scaffold00](http://itak.feilab.net/cgi-bin/itak/db_gene_seq.cgi?trans_ID=evm_27.model.AmTr_v1.0_scaffold00127.17)  [127.17](http://itak.feilab.net/cgi-bin/itak/db_gene_seq.cgi?trans_ID=evm_27.model.AmTr_v1.0_scaffold00127.17) | [ERM97390.1](https://www.ncbi.nlm.nih.gov/protein/ERM97390.1) | [AMTR_s001](https://alphafold.ebi.ac.uk/search/text/AMTR_s00127p00060060) [27p0006006](https://alphafold.ebi.ac.uk/search/text/AMTR_s00127p00060060)  [0](https://alphafold.ebi.ac.uk/search/text/AMTR_s00127p00060060) | [W1NQY5](https://www.uniprot.org/uniprotkb/W1NQY5/entry) | AmSVP |
| SVP | *Vitis vinifera* | Uncharacterized protein | [GSVIVT010](http://itak.feilab.net/cgi-bin/itak/db_gene_seq.cgi?trans_ID=GSVIVT01009219001) [09219001](http://itak.feilab.net/cgi-bin/itak/db_gene_seq.cgi?trans_ID=GSVIVT01009219001) | [CBI19301.3](https://www.ncbi.nlm.nih.gov/protein/CBI19301.3) | [VIT_18s000](https://alphafold.ebi.ac.uk/search/text/VIT_18s0001g07900) [1g07900](https://alphafold.ebi.ac.uk/search/text/VIT_18s0001g07900) | [E0CNS3](https://www.uniprot.org/uniprotkb/E0CNS3/entry) | VviAGL16 |
| SVP | *Vitis vinifera* | Uncharacterized protein | [GSVIVT010](http://itak.feilab.net/cgi-bin/itak/db_gene_seq.cgi?trans_ID=GSVIVT01003864001) [03864001](http://itak.feilab.net/cgi-bin/itak/db_gene_seq.cgi?trans_ID=GSVIVT01003864001) | [XP_0022735](https://www.ncbi.nlm.nih.gov/protein/XP_002273556.2) [56.2](https://www.ncbi.nlm.nih.gov/protein/XP_002273556.2) | [VIT_00s021](https://alphafold.ebi.ac.uk/search/text/VIT_00s0211g00180) [1g00180](https://alphafold.ebi.ac.uk/search/text/VIT_00s0211g00180) | [D7T5C5](https://www.uniprot.org/uniprotkb/D7T5C5/entry) | VviAGL21 |

| **Clade** | **Organism** | **Protein name** | **iTAK, TAIR**  **or PlanTFDB Identifier** | **NCBI**  **Identifier** | **AlphaFold Identifier** | **Identifier Uniprot** | **Gene name** |
| --- | --- | --- | --- | --- | --- | --- | --- |
| SVP | *Arabidopsis thaliana* | Agamous-like MADS-box protein AGL17 | [AT2G22630.](https://www.arabidopsis.org/servlets/TairObject?id=35530&type=locus)  [1](https://www.arabidopsis.org/servlets/TairObject?id=35530&type=locus) | [NP_0013244](https://www.ncbi.nlm.nih.gov/protein/NP_001324454.1) [54.1](https://www.ncbi.nlm.nih.gov/protein/NP_001324454.1) | [AGL17_ARA](https://alphafold.ebi.ac.uk/search/text/AGL17_ARATH) [TH](https://alphafold.ebi.ac.uk/search/text/AGL17_ARATH) | [Q38840](https://www.uniprot.org/uniprotkb/Q38840/entry) | AtAGL17 |
| SVP | *Arabidopsis thaliana* | Agamous-like MADS-box protein AGL21 | [AT4G37940.](https://www.arabidopsis.org/servlets/TairObject?id=127455&type=locus)  [1](https://www.arabidopsis.org/servlets/TairObject?id=127455&type=locus) | [NP_195507.](https://www.ncbi.nlm.nih.gov/protein/NP_195507.1)  [1](https://www.ncbi.nlm.nih.gov/protein/NP_195507.1) | [AGL21_ARA](https://alphafold.ebi.ac.uk/search/text/AGL21_ARATH) [TH](https://alphafold.ebi.ac.uk/search/text/AGL21_ARATH) | [Q9SZJ6](https://www.uniprot.org/uniprotkb/Q9SZJ6/entry) | AtAGL21 |
| SVP | *Arabidopsis thaliana* | Agamous-like MADS-box protein AGL16 | [AT3G57230.](https://www.arabidopsis.org/servlets/TairObject?id=37044&type=locus)  [1](https://www.arabidopsis.org/servlets/TairObject?id=37044&type=locus) | [NP_0013262](https://www.ncbi.nlm.nih.gov/protein/NP_001326287.1) [87.1](https://www.ncbi.nlm.nih.gov/protein/NP_001326287.1) | [AGL16_ARA](https://alphafold.ebi.ac.uk/search/text/AGL16_ARATH) [TH](https://alphafold.ebi.ac.uk/search/text/AGL16_ARATH) | [A2RVQ5](https://www.uniprot.org/uniprotkb/A2RVQ5/entry) | AtAGL16 |
| SVP | *Arabidopsis thaliana* | AGAMOUS-like 44 | [AT2G14210.](https://www.arabidopsis.org/servlets/TairObject?id=31853&type=locus)  [2](https://www.arabidopsis.org/servlets/TairObject?id=31853&type=locus) | [NP_0013240](https://www.ncbi.nlm.nih.gov/protein/NP_001324071.1) [71.1](https://www.ncbi.nlm.nih.gov/protein/NP_001324071.1) | [A0A1P8AYX](https://alphafold.ebi.ac.uk/search/text/A0A1P8AYX0_ARATH) [0_ARATH](https://alphafold.ebi.ac.uk/search/text/A0A1P8AYX0_ARATH) | [A0A1P8AYX](https://www.uniprot.org/uniprotkb/A0A1P8AYX0/entry) [0](https://www.uniprot.org/uniprotkb/A0A1P8AYX0/entry) | AtAGL44 |
| SVP | *Amborella trichopoda* | Uncharacterized protein | [evm_27.mod](http://itak.feilab.net/cgi-bin/itak/db_gene_seq.cgi?trans_ID=evm_27.model.AmTr_v1.0_scaffold00046.134) [el.AmTr_v1.](http://itak.feilab.net/cgi-bin/itak/db_gene_seq.cgi?trans_ID=evm_27.model.AmTr_v1.0_scaffold00046.134) [0_scaffold00](http://itak.feilab.net/cgi-bin/itak/db_gene_seq.cgi?trans_ID=evm_27.model.AmTr_v1.0_scaffold00046.134)  [046.134](http://itak.feilab.net/cgi-bin/itak/db_gene_seq.cgi?trans_ID=evm_27.model.AmTr_v1.0_scaffold00046.134) | [ERN18058.1](https://www.ncbi.nlm.nih.gov/protein/ERN18058.1) | [AMTR_s000](https://alphafold.ebi.ac.uk/search/text/AMTR_s00046p00208680) [46p0020868](https://alphafold.ebi.ac.uk/search/text/AMTR_s00046p00208680)  [0](https://alphafold.ebi.ac.uk/search/text/AMTR_s00046p00208680) | [U5DCD9](https://www.uniprot.org/uniprotkb/U5DCD9/entry) | AmAGL21 |
| MIKC* | *Marchantia polymorpha* | MIKC* MADS-  box transcription factor | [Mapoly0174](http://planttfdb.gao-lab.org/tf.php?sp=Mpo&did=Mapoly0174s0011.1.p) [s0011.1.p](http://planttfdb.gao-lab.org/tf.php?sp=Mpo&did=Mapoly0174s0011.1.p) | [ADB81895.1](https://www.ncbi.nlm.nih.gov/protein/ADB81895.1) | [D3IZT6_MA](https://alphafold.ebi.ac.uk/search/text/D3IZT6_MARPO) [RPO](https://alphafold.ebi.ac.uk/search/text/D3IZT6_MARPO) | [D3IZT6](https://www.uniprot.org/uniprotkb/D3IZT6/entry) | MpMADS1 |
| MIKC* | *Arabidopsis thaliana* | Agamous-like MADS-box  protein AGL63 | [AT1G31140.](https://www.arabidopsis.org/servlets/TairObject?id=29827&type=locus)  [2](https://www.arabidopsis.org/servlets/TairObject?id=29827&type=locus) | [NP_0011851](https://www.ncbi.nlm.nih.gov/protein/NP_001185120.1) [20.1](https://www.ncbi.nlm.nih.gov/protein/NP_001185120.1) | [AGL63_ARA](https://alphafold.ebi.ac.uk/search/text/AGL63_ARATH) [TH](https://alphafold.ebi.ac.uk/search/text/AGL63_ARATH) | [Q9SA07](https://www.uniprot.org/uniprotkb/Q9SA07/entry) | AtGOA |
| MIKC* | *Arabidopsis thaliana* | MADS-box domain-containing  protein | [AT1G77950.](https://www.arabidopsis.org/servlets/TairObject?id=29803&type=locus)  [1](https://www.arabidopsis.org/servlets/TairObject?id=29803&type=locus) | [NP_0011176](https://www.ncbi.nlm.nih.gov/protein/NP_001117616.1?report=genbank&log%24=prottop&blast_rank=1&RID=V9PWJW72013) [16.1](https://www.ncbi.nlm.nih.gov/protein/NP_001117616.1?report=genbank&log%24=prottop&blast_rank=1&RID=V9PWJW72013) | [A0A654EV2](https://alphafold.ebi.ac.uk/search/text/A0A654EV27) [7_ARATH](https://alphafold.ebi.ac.uk/search/text/A0A654EV27) | [A0A654EV2](https://www.uniprot.org/uniprotkb/A0A654EV27/entry) [7](https://www.uniprot.org/uniprotkb/A0A654EV27/entry) | AtAGL67 |
| MIKC* | *Arabidopsis thaliana* | MADS-box domain-  containing protein | [AT1G77980.](https://www.arabidopsis.org/servlets/TairObject?id=29798&type=locus)  [1](https://www.arabidopsis.org/servlets/TairObject?id=29798&type=locus) | [NP_177921.](https://www.ncbi.nlm.nih.gov/protein/NP_177921.2?report=genbank&log%24=prottop&blast_rank=1&RID=V9PWJW72013)  [2](https://www.ncbi.nlm.nih.gov/protein/NP_177921.2?report=genbank&log%24=prottop&blast_rank=1&RID=V9PWJW72013) | [A0A654EQ6](https://alphafold.ebi.ac.uk/search/text/A0A654EQ65) [5_ARATH](https://alphafold.ebi.ac.uk/search/text/A0A654EQ65) | [A0A654EQ6](https://www.uniprot.org/uniprotkb/A0A654EQ65/entry) [5](https://www.uniprot.org/uniprotkb/A0A654EQ65/entry) | AtAGL66 |
| MIKC* | *Arabidopsis thaliana* | MADS-box domain-containing  protein | [AT1G22130.](https://www.arabidopsis.org/servlets/TairObject?id=29996&type=locus)  [1](https://www.arabidopsis.org/servlets/TairObject?id=29996&type=locus) | [NP_173632.](https://www.ncbi.nlm.nih.gov/protein/NP_173632.1?report=genbank&log%24=prottop&blast_rank=1&RID=V9PWJW72013)  [1](https://www.ncbi.nlm.nih.gov/protein/NP_173632.1?report=genbank&log%24=prottop&blast_rank=1&RID=V9PWJW72013) | [A0A654EBX](https://alphafold.ebi.ac.uk/search/text/A0A654EBX2) [2_ARATH](https://alphafold.ebi.ac.uk/search/text/A0A654EBX2) | [A0A654EBX](https://www.uniprot.org/uniprotkb/A0A654EBX2/entry) [2](https://www.uniprot.org/uniprotkb/A0A654EBX2/entry) | AtAGL104 |
| MIKC* | *Arabidopsis thaliana* | AGAMOUS-like 65 | [AT1G18750.](https://www.arabidopsis.org/servlets/TairObject?id=30666&type=locus)  [3](https://www.arabidopsis.org/servlets/TairObject?id=30666&type=locus) | [NP_0013215](https://www.ncbi.nlm.nih.gov/protein/NP_001321504.1?report=genbank&log%24=prottop&blast_rank=1&RID=V9PWJW72013) [04.1](https://www.ncbi.nlm.nih.gov/protein/NP_001321504.1?report=genbank&log%24=prottop&blast_rank=1&RID=V9PWJW72013) | [A0A1P8AR6](https://alphafold.ebi.ac.uk/search/text/A0A1P8AR61) [1_ARATH](https://alphafold.ebi.ac.uk/search/text/A0A1P8AR61) | [A0A1P8AR6](https://www.uniprot.org/uniprotkb/A0A1P8AR61/entry#sequences) [1](https://www.uniprot.org/uniprotkb/A0A1P8AR61/entry#sequences) | AtAGL65 |
| MIKC* | *Arabidopsis thaliana* | AGAMOUS-like 94 | [AT1G69540.](https://www.arabidopsis.org/servlets/TairObject?id=26681&type=locus)  [2](https://www.arabidopsis.org/servlets/TairObject?id=26681&type=locus) | [NP_0013221](https://www.ncbi.nlm.nih.gov/protein/NP_001322116.1?report=genbank&log%24=prottop&blast_rank=1&RID=V9PWJW72013) [16.1](https://www.ncbi.nlm.nih.gov/protein/NP_001322116.1?report=genbank&log%24=prottop&blast_rank=1&RID=V9PWJW72013) | [A0A1P8ASZ](https://alphafold.ebi.ac.uk/search/text/A0A1P8ASZ9) [9_ARATH](https://alphafold.ebi.ac.uk/search/text/A0A1P8ASZ9) | [A0A1P8ASZ](https://www.uniprot.org/uniprotkb/A0A1P8ASZ9/entry) [9](https://www.uniprot.org/uniprotkb/A0A1P8ASZ9/entry) | AtAGL94 |
| MIKC* | *Arabidopsis thaliana* | Agamous-like  MADS-box protein AGL30 | [AT2G03060.](https://www.arabidopsis.org/servlets/TairObject?id=34056&type=locus)  [2](https://www.arabidopsis.org/servlets/TairObject?id=34056&type=locus) | [NP_0013181](https://www.ncbi.nlm.nih.gov/protein/NP_001318187.1?report=genbank&log%24=prottop&blast_rank=1&RID=V9PWJW72013) [87.1](https://www.ncbi.nlm.nih.gov/protein/NP_001318187.1?report=genbank&log%24=prottop&blast_rank=1&RID=V9PWJW72013) | [AGL30_ARA](https://alphafold.ebi.ac.uk/search/text/Q1PFA4) [TH](https://alphafold.ebi.ac.uk/search/text/Q1PFA4) | [Q1PFA4-1](https://www.uniprot.org/uniprotkb/Q1PFA4/entry#Q1PFA4-1) | AtAGL30 |
| TT16 | *Vitis vinifera* | Uncharacterized protein | - | [XP_0022719](https://www.ncbi.nlm.nih.gov/protein/XP_002271905.2) [05.2](https://www.ncbi.nlm.nih.gov/protein/XP_002271905.2) | [VIT_10s004](https://alphafold.ebi.ac.uk/search/text/VIT_10s0042g00820) [2g00820](https://alphafold.ebi.ac.uk/search/text/VIT_10s0042g00820) | [F6HIR2](https://www.uniprot.org/uniprotkb/F6HIR2/entry) | VviTT16a |
| TT16 | *Arabidopsis thaliana* | K-box region and MADS-box transcription  factor family protein | [AT5G23260.](https://www.arabidopsis.org/servlets/TairObject?id=133655&type=locus)  [4](https://www.arabidopsis.org/servlets/TairObject?id=133655&type=locus) | [NP_0013304](https://www.ncbi.nlm.nih.gov/protein/NP_001330404.1) [04.1](https://www.ncbi.nlm.nih.gov/protein/NP_001330404.1) | [A0A1P8BA](https://alphafold.ebi.ac.uk/search/text/A0A1P8BAR2_ARATH) [R2_ARATH](https://alphafold.ebi.ac.uk/search/text/A0A1P8BAR2_ARATH) | [A0A1P8BAR](https://www.uniprot.org/uniprotkb/A0A1P8BAR2/entry) [2](https://www.uniprot.org/uniprotkb/A0A1P8BAR2/entry) | AtTT16 |
| TT16 | *Vitis vinifera* | Uncharacterized protein | - | [XP_0190767](https://www.ncbi.nlm.nih.gov/protein/XP_019076763.1) [63.1](https://www.ncbi.nlm.nih.gov/protein/XP_019076763.1)  ** | [VIT_01s001](https://alphafold.ebi.ac.uk/search/text/VIT_01s0011g01560) [1g01560](https://alphafold.ebi.ac.uk/search/text/VIT_01s0011g01560) | [F6HF63](https://www.uniprot.org/uniprotkb/F6HF63/entry) | VviTT16b |
| TT16 | *Vitis vinifera* | Uncharacterized protein | - | [XP_0106598](https://www.ncbi.nlm.nih.gov/protein/XP_010659881.1) [81.1](https://www.ncbi.nlm.nih.gov/protein/XP_010659881.1)  ** | [VIT_02s002](https://alphafold.ebi.ac.uk/search/text/VIT_02s0025g02350) [5g02350](https://alphafold.ebi.ac.uk/search/text/VIT_02s0025g02350) | [F6HU63](https://www.uniprot.org/uniprotkb/F6HU63/entry) | VviTT16c |

| **Clade** | **Organism** | **Protein name** | **iTAK, TAIR**  **or PlanTFDB Identifier** | **NCBI**  **Identifier** | **AlphaFold Identifier** | **Identifier Uniprot** | **Gene name** |
| --- | --- | --- | --- | --- | --- | --- | --- |
| TT16 | Amborella trichopoda | Uncharacterized protein | [evm_27.mod](http://itak.feilab.net/cgi-bin/itak/db_gene_seq.cgi?trans_ID=evm_27.model.AmTr_v1.0_scaffold00001.461) [el.AmTr_v1.](http://itak.feilab.net/cgi-bin/itak/db_gene_seq.cgi?trans_ID=evm_27.model.AmTr_v1.0_scaffold00001.461) [0_scaffold](http://itak.feilab.net/cgi-bin/itak/db_gene_seq.cgi?trans_ID=evm_27.model.AmTr_v1.0_scaffold00001.461)  [00001.461](http://itak.feilab.net/cgi-bin/itak/db_gene_seq.cgi?trans_ID=evm_27.model.AmTr_v1.0_scaffold00001.461) | [ERM96584.1](https://www.ncbi.nlm.nih.gov/protein/ERM96584.1) | [AMTR_s000](https://alphafold.ebi.ac.uk/search/text/AMTR_s00001p00270400) [01p0027040](https://alphafold.ebi.ac.uk/search/text/AMTR_s00001p00270400)  [0](https://alphafold.ebi.ac.uk/search/text/AMTR_s00001p00270400) | [W1NMR3](https://www.uniprot.org/uniprotkb/W1NMR3/entry) | AmTT16 |
| Pistillata | *Vitis vinifera* | Agamous-like MADS-box protein MADS9 | - | [Q0HA25.2](https://www.ncbi.nlm.nih.gov/protein/Q0HA25.2) | [VvPI](https://alphafold.ebi.ac.uk/search/text/VvPI) | [Q0HA25](https://www.uniprot.org/uniprotkb/Q0HA25/entry) | VviPI |
| Pistillata | *Arabidopsis thaliana* | Floral homeotic protein PISTILLATA | [AT5G20240.](https://www.arabidopsis.org/servlets/TairObject?id=131273&type=locus)  [1](https://www.arabidopsis.org/servlets/TairObject?id=131273&type=locus) | [NP_197524.](https://www.ncbi.nlm.nih.gov/protein/NP_197524.1)  [1](https://www.ncbi.nlm.nih.gov/protein/NP_197524.1) | [PIST_ARAT](https://alphafold.ebi.ac.uk/search/text/PIST_ARATH) [H](https://alphafold.ebi.ac.uk/search/text/PIST_ARATH) | [P48007](https://www.uniprot.org/uniprotkb/P48007/entry) | AtPI |
| Pistillata | *Amborella trichopoda* | PISTILLATA-like  protein | - | [BAD42443.1](https://www.ncbi.nlm.nih.gov/protein/BAD42443.1) | [Q68BI0_AM](https://alphafold.ebi.ac.uk/search/text/Q68BI0_AMBTC) [BTC](https://alphafold.ebi.ac.uk/search/text/Q68BI0_AMBTC) | [Q68BI0](https://www.uniprot.org/uniprotkb/Q68BI0/entry) | AmPIa |
| Pistillata | *Amborella trichopoda* | Uncharacterized protein | [evm_27.mod](http://itak.feilab.net/cgi-bin/itak/db_gene_seq.cgi?trans_ID=evm_27.model.AmTr_v1.0_scaffold00089.36) [el.AmTr_v1.](http://itak.feilab.net/cgi-bin/itak/db_gene_seq.cgi?trans_ID=evm_27.model.AmTr_v1.0_scaffold00089.36) [0_scaffold](http://itak.feilab.net/cgi-bin/itak/db_gene_seq.cgi?trans_ID=evm_27.model.AmTr_v1.0_scaffold00089.36)  [00089.36](http://itak.feilab.net/cgi-bin/itak/db_gene_seq.cgi?trans_ID=evm_27.model.AmTr_v1.0_scaffold00089.36) | [XP_0068401](https://www.ncbi.nlm.nih.gov/protein/XP_006840164.1) [64.1](https://www.ncbi.nlm.nih.gov/protein/XP_006840164.1) | [AMTR_s000](https://alphafold.ebi.ac.uk/search/text/AMTR_s00089p00081270) [89p0008127](https://alphafold.ebi.ac.uk/search/text/AMTR_s00089p00081270)  [0](https://alphafold.ebi.ac.uk/search/text/AMTR_s00089p00081270) | [W1P2H0](https://www.uniprot.org/uniprotkb/W1P2H0/entry) | AmPIb |
| AP3 | *Amborella trichopoda* | Uncharacterized protein | [evm_27.](http://itak.feilab.net/cgi-bin/itak/db_gene_seq.cgi?trans_ID=evm_27.model.AmTr_v1.0_scaffold00066.97) [model.AmTr](http://itak.feilab.net/cgi-bin/itak/db_gene_seq.cgi?trans_ID=evm_27.model.AmTr_v1.0_scaffold00066.97)  [_v1.0_scaffo](http://itak.feilab.net/cgi-bin/itak/db_gene_seq.cgi?trans_ID=evm_27.model.AmTr_v1.0_scaffold00066.97) [ld00066.97](http://itak.feilab.net/cgi-bin/itak/db_gene_seq.cgi?trans_ID=evm_27.model.AmTr_v1.0_scaffold00066.97) | [XP_0068587](https://www.ncbi.nlm.nih.gov/protein/XP_006858714.1) [14.1](https://www.ncbi.nlm.nih.gov/protein/XP_006858714.1) | [AMTR_s000](https://alphafold.ebi.ac.uk/search/text/AMTR_s00066p00109320) [66p0010932](https://alphafold.ebi.ac.uk/search/text/AMTR_s00066p00109320)  [0](https://alphafold.ebi.ac.uk/search/text/AMTR_s00066p00109320) | [U5D3L0](https://www.uniprot.org/uniprotkb/U5D3L0/entry) | AmAP3a |
| AP3 | *Amborella trichopoda* | APETALA3-like protein | - | [BAD42444.1](https://www.ncbi.nlm.nih.gov/protein/BAD42444.1) | [Q68BH9_A](https://alphafold.ebi.ac.uk/search/text/Q68BH9_AMBTC) [MBTC](https://alphafold.ebi.ac.uk/search/text/Q68BH9_AMBTC) | [Q68BH9](https://www.uniprot.org/uniprotkb/Q68BH9/entry) | AmAP3b |
| AP3 | *Amborella trichopoda* | APETALA3-like protein AP3-1 | - | [AAR06677.1](https://www.ncbi.nlm.nih.gov/protein/AAR06677.1) | [I6LAR5_AM](https://alphafold.ebi.ac.uk/search/text/I6LAR5_AMBTC) [BTC](https://alphafold.ebi.ac.uk/search/text/I6LAR5_AMBTC) | [I6LAR5](https://www.uniprot.org/uniprotkb/I6LAR5/entry) | AmAP3c |
| AP3 | *Amborella trichopoda* | Uncharacterized protein | [evm_27.mod](http://itak.feilab.net/cgi-bin/itak/db_gene_seq.cgi?trans_ID=evm_27.model.AmTr_v1.0_scaffold00001.225) [el.AmTr_v1.](http://itak.feilab.net/cgi-bin/itak/db_gene_seq.cgi?trans_ID=evm_27.model.AmTr_v1.0_scaffold00001.225) [0_scaffold](http://itak.feilab.net/cgi-bin/itak/db_gene_seq.cgi?trans_ID=evm_27.model.AmTr_v1.0_scaffold00001.225)  [00001.225](http://itak.feilab.net/cgi-bin/itak/db_gene_seq.cgi?trans_ID=evm_27.model.AmTr_v1.0_scaffold00001.225) | [ERM96348.1](https://www.ncbi.nlm.nih.gov/protein/ERM96348.1) | [AMTR_s000](https://alphafold.ebi.ac.uk/search/text/AMTR_s00001p00217560) [01p0021756](https://alphafold.ebi.ac.uk/search/text/AMTR_s00001p00217560)  [0](https://alphafold.ebi.ac.uk/search/text/AMTR_s00001p00217560) | [W1NLE9](https://www.uniprot.org/uniprotkb/W1NLE9/entry) | AmAP3d |
| AP3 | Vitis vinifera | Agamous-like MADS-box  protein AP3 | [GSVIVT010](http://itak.feilab.net/cgi-bin/itak/db_gene_seq.cgi?trans_ID=GSVIVT01009815001) [09815001](http://itak.feilab.net/cgi-bin/itak/db_gene_seq.cgi?trans_ID=GSVIVT01009815001) | [E0CPH4.1](https://www.ncbi.nlm.nih.gov/protein/E0CPH4.1) | [AP3_VITVI](https://alphafold.ebi.ac.uk/search/text/AP3_VITVI) | [E0CPH4](https://www.uniprot.org/uniprotkb/E0CPH4/entry) | VviAP3 |
| AP3 | *Arabidopsis thaliana* | Floral homeotic protein APETALA 3 | [AT3G54340.](https://www.arabidopsis.org/servlets/TairObject?id=39428&type=locus)  [1](https://www.arabidopsis.org/servlets/TairObject?id=39428&type=locus) | [NP_191002.](https://www.ncbi.nlm.nih.gov/protein/NP_191002.1)  [1](https://www.ncbi.nlm.nih.gov/protein/NP_191002.1) | [AP3_ARATH](https://alphafold.ebi.ac.uk/search/text/AP3_ARATH) | [P35632](https://www.uniprot.org/uniprotkb/P35632/entry) | AtAP3 |
| FLC | *Arabidopsis thaliana* | Agamous-like MADS-box protein AGL70 | [AT5G65060.](https://www.arabidopsis.org/servlets/TairObject?id=130633&type=locus)  [1](https://www.arabidopsis.org/servlets/TairObject?id=130633&type=locus) | [NP_201311.](https://www.ncbi.nlm.nih.gov/protein/NP_201311.1)  [1](https://www.ncbi.nlm.nih.gov/protein/NP_201311.1) | [AGL70_ARA](https://alphafold.ebi.ac.uk/search/text/AGL70_ARATH) [TH](https://alphafold.ebi.ac.uk/search/text/AGL70_ARATH) | [Q9LSR7](https://www.uniprot.org/uniprotkb/Q9LSR7/entry) | AtMAF3 |
| FLC | *Arabidopsis thaliana* | Agamous-like MADS-box protein AGL31 | [AT5G65050.](https://www.arabidopsis.org/servlets/TairObject?id=135154&type=locus)  [3](https://www.arabidopsis.org/servlets/TairObject?id=135154&type=locus) | [NP_0011194](https://www.ncbi.nlm.nih.gov/protein/NP_001119498.1) [98.1](https://www.ncbi.nlm.nih.gov/protein/NP_001119498.1) | [AGL31_ARA](https://alphafold.ebi.ac.uk/search/text/AGL31_ARATH) [TH](https://alphafold.ebi.ac.uk/search/text/AGL31_ARATH) | [Q9FPN7](https://www.uniprot.org/uniprotkb/Q9FPN7/entry) | AtMAF2 |
| FLC | *Arabidopsis thaliana* | K-box region and MADS-box transcription factor family protein | [AT1G77080.](https://www.arabidopsis.org/servlets/TairObject?id=29106&type=locus)  [8](https://www.arabidopsis.org/servlets/TairObject?id=29106&type=locus) | [NP_0013215](https://www.ncbi.nlm.nih.gov/protein/NP_001321570.1) [70.1](https://www.ncbi.nlm.nih.gov/protein/NP_001321570.1) | [A0A1P8ARA](https://alphafold.ebi.ac.uk/search/text/A0A1P8ARA3_ARATH) [3_ARATH](https://alphafold.ebi.ac.uk/search/text/A0A1P8ARA3_ARATH) | [A0A1P8ARA](https://www.uniprot.org/uniprotkb/A0A1P8ARA3/entry) [3](https://www.uniprot.org/uniprotkb/A0A1P8ARA3/entry) | AtMAF1 |
| FLC | *Arabidopsis thaliana* | K-box region and MADS-box transcription factor family protein | [AT5G65070.](https://www.arabidopsis.org/servlets/TairObject?id=130634&type=locus)  [3](https://www.arabidopsis.org/servlets/TairObject?id=130634&type=locus) | [NP_0011906](https://www.ncbi.nlm.nih.gov/protein/NP_001190617.1) [17.1](https://www.ncbi.nlm.nih.gov/protein/NP_001190617.1) | [F4KGH9_AR](https://alphafold.ebi.ac.uk/search/text/F4KGH9_ARATH) [ATH](https://alphafold.ebi.ac.uk/search/text/F4KGH9_ARATH) | [F4KGH9](https://www.uniprot.org/uniprotkb/F4KGH9/entry) | AtMAF4 |

| **Clade** | **Organism** | **Protein name** | **iTAK, TAIR**  **or PlanTFDB Identifier** | **NCBI**  **Identifier** | **AlphaFold Identifier** | **Identifier Uniprot** | **Gene name** |
| --- | --- | --- | --- | --- | --- | --- | --- |
| FLC | *Arabidopsis thaliana* | K-box region/MADS-box transcription factor family  protein | [AT5G65080.](https://www.arabidopsis.org/servlets/TairObject?id=130635&type=locus)  [2](https://www.arabidopsis.org/servlets/TairObject?id=130635&type=locus) | [NP_0010787](https://www.ncbi.nlm.nih.gov/protein/NP_001078799.2) [99.2](https://www.ncbi.nlm.nih.gov/protein/NP_001078799.2) | [A0A2H1ZE](https://alphafold.ebi.ac.uk/search/text/A0A2H1ZE96_ARATH) [96_ARATH](https://alphafold.ebi.ac.uk/search/text/A0A2H1ZE96_ARATH) | [A0A2H1ZE9](https://www.uniprot.org/uniprotkb/A0A2H1ZE96/entry) [6](https://www.uniprot.org/uniprotkb/A0A2H1ZE96/entry) | AtMAF5 |
| FLC | *Arabidopsis thaliana* | MADS-box protein FLOWERING  LOCUS C | [AT5G10140.](https://www.arabidopsis.org/servlets/TairObject?id=136002&type=locus)  [1](https://www.arabidopsis.org/servlets/TairObject?id=136002&type=locus) | [NP_196576.](https://www.ncbi.nlm.nih.gov/protein/NP_196576.1)  [1](https://www.ncbi.nlm.nih.gov/protein/NP_196576.1) | [FLC_ARATH](https://alphafold.ebi.ac.uk/search/text/FLC_ARATH) | [Q9S7Q7](https://www.uniprot.org/uniprotkb/Q9S7Q7/entry) | AtFLC |
